# Impact of force function formulations on the numerical simulation of centre-based models

**DOI:** 10.1101/2020.03.16.993246

**Authors:** Sonja Mathias, Adrien Coulier, Anass Bouchnita, Andreas Hellander

## Abstract

Centre-based, or cell-centre models are a framework for the computational study of multicellular systems with widespread use in cancer modelling and computational developmental biology. At the core of these models are the numerical method used to update cell positions and the force functions that encode the pairwise mechanical interactions of cells. For the latter there are multiple choices that could potentially affect both the biological behaviour captured, and the robustness and efficiency of simulation. For example, available open-source software implementations of centre-based models rely on different force functions for their default behaviour and it is not straightforward for a modeler to know if these are interchangeable. Our study addresses this problem and contributes to the understanding of the potential and limitations of three popular force functions from a numerical perspective. We show empirically that choosing the force parameters such that the relaxation time for two cells after cell division is consistent between different force functions results in good agreement of the population radius of a growing monolayer. Furthermore, we report that numerical stability is not sufficient to prevent unphysical cell trajectories following cell division, and consequently, that too large time steps can cause geometrical differences at the population level.

## 1 Introduction

Discrete cell-based models are becoming increasingly popular for simulating tissue mechanics. In contrast to continuum models that average over the cell density, cell-based models represent each cell individually. Therefore, they readily allow for incorporating cellular events such as cell division, cell differentiation or cell death, as well as cell heterogeneity across a population. As a result, these models have been used to probe biological questions of how the interplay of individual cell behaviour affects population-level measures (e.g. [1, 2, 3, 4]).

There are different kinds of cell-based models, that can be divided into two general categories. The first category, so-called *on-lattice* models, restricts the movement of the cells to a grid. Cellular automata [5] and cellular Potts [6] models are examples. In cellular automata models, cells are typically restricted to occupy a single lattice site and move between lattice sites according to a fixed set of rules. In contrast, in cellular Potts models cells are composed of multiple lattice sites, enabling the cell shape to be resolved more realistically. The whole system explores the energy landscape using a Metropolis-Hastings approach. One drawback of on-lattice models is that they can exhibit grid-related artefacts on structured meshes due to the directional restriction, e.g. cells can only “push” neighbours along fixed axes as defined by the underlying grid [7, 8].

The second category, *off-lattice* models, are continuous in space and hence circumvent this issue. Again they vary with respect to how detailed the cell shape is modelled. Centre-based models (CBMs) — also referred to as cell-centre models — track the cell midpoints over time as cells interact mechanically according to pairwise spring-like forces [1, 9]. In this model, cells are either represented as overlapping spheres (OS variant), or using a Voronoi tessellation (Voronoi variant). Vertex models [10], on the other hand, discretize the cell boundary instead and evolve the tissue according to interfacial tension and pressure within the cells. At an even higher level of detail and correspondingly higher computational cost there are the immersed boundary method [11] and the subcellular element method [12].

Discrete cell-based models — independent of being on- or off-lattice — can be coupled to PDE models for simulating the concentration of chemical compounds in the cellular environment or even an ODE model for simulating intracellular dynamics [13, 14, 15]. An extensive review of cell-based models for general tissue mechanics can be found in [7]. Additionally, there are several reviews dealing with prominent applications areas, such as tumour growth [16, 17] and morphogenetic problems [18, 19, 20]. In [21] the authors compare five cell-based frameworks (cellular automata, cellular potts, CBM OS and Voronoi variants and vertex models) with respect to four common biological problems: cell sorting, monoclonal conversion, lateral inihibition and morphogen-dependent proliferation. They conclude that each model has its preferred application for the study of which it was originally designed, but that most models can be adapted for all applications with varying effort and computational cost.

In this study we focus on the centre-based model, in particular the OS variant, to which we will from now on refer to as CBM or CBM OS when we want to stress particularities about the latter. CBMs have been successfully applied to a large variety of biological problems ranging from the simulation of monolayer and spheroid growth [9, 22] to the cellular reorganization in the intestinal crypt [1]. See [23] for a recent overview. There exist multiple simulation frameworks that implement CBMs, several of which are open-source. All of them tailor to specific needs, but allow for modelling the core features of CBMs. *Chaste* is a multi-purpose framework implementing several cell-based models and CBMs in particular [24, 25]. *MecaGen* is a framework focusing on the coupling between cell mechanics and gene regulatory networks [26]. Most recently, *PhysiCell* was released in 2018 [27]. It aims to simulate up to a million cells and has been used mainly to model breast cancer [28, 29, 30]. Moreover, there exist the closed-source frameworks *CellSys* [31] and *Biocellion* [32].

In general, one can observe the trend of simulating an ever-growing number of cells in order to realistically describe biological systems. The efficiency of simulations is critical to this endeavor. Highly efficient schemes not only allow for the simulation of large-scale systems, but also make systematic inference and model exploration feasible. Here, the CBM framework, and in particular the OS variant in its simplest form, strikes an attractive balance between numerical efficiency and the capability to incorporate cell mechanics and biophysical measurements. The biophysical basis of cell-cell interactions has been discussed extensively in prior work [9, 33, 34], where the Johnson-Kendall-Roberts (JKR) and extended Hertz theories can provide descriptions founded in physics [35]. However, these force descriptions are computationally expensive to evaluate, and in practice other, simpler functional forms are considered.

Motivated by the need for robust and efficient simulations, this study focuses on the complementary question of comparing those force function used in open-source frameworks for CBMs. More specifically, we choose the functions used in Chaste, PhysiCell and MecaGen [24, 27, 26]. Given the relatively few numbers of free parameters in these force functions, we first ask if there are systematic ways to parameterize them such that they result in indistinguishable macroscopic output for a biologically realistic test problem. We show that it is sufficient to accurately match the relaxation time of cells after division in order to observe good agreement of trajectories for the population radius of a freely growing monolayer. Based on such a parameterization, we then ask if there are significant advantages and disadvantages of different force functions from a numerical robustness- and efficiency point of view.

Different solution strategies and integration schemes for Vertex Models were discussed and evaluated in detail in [36]. However, to the best of our knowledge a systematic comparison of different commonly used schemes for CBMs has not been published. The overwhelming majority of CBM implementations use the Euler forward method for solving the coupled ODE system for centre positions [24, 26, 37], with PhysiCell being a notable exception in that it uses the second-order Adams-Bashford method [27]. [38] study mechanical properties of a tissue simulated with CBMs — both the OS and the Voronoi variants — subject to compression, tension and shear under a linear spring force and two physics-based forces, Hertz and Johnson-Kendall-Roberts. Instead of probing the mechanical properties of a non-proliferating tissue, we are concerned with the interplay between cell proliferation and the impact of the resulting high local forces on the mechanical relaxation of a growing population. Here, a critical aspect is the potentially large stiffness caused by highly compressed cells right after cell division. A key contribution of this paper is a comparison of integration schemes for this setting, taking into account the dependence on the force functions for stability and error.

This article is structured as follows. In Section 2 we review the key numerical aspects of CBMs. In Section 3, we describe three force functions encountered in popular simulation frameworks and compare their qualitative behaviour for a fixed parameterization. In Section 4 we study their numerical properties with regards to numerical stability and accuracy in combination with first- and second-order solvers. Lastly, we discuss our findings and conclusions in Section 5.

## 2 Implementation of centre-based models

In this section, we review the key numerical assumptions made when implementing CBMs, namely (i) the update equation for cell positions, (ii) the numerical solution of this equation, (iii) the way cell connectivity is defined and updated, (iv) how cell-level events are incorporated into the cell mechanics simulation and (v) how the cell cycle and cell division events are modelled in particular. In literature and existing software, there is a considerable variation when it comes to the details of the implementations, depending on the biophysiology studied and the computational budget. Motivated by the need to simulate increasingly large systems, we consider the simplest and computationally cheapest implementations as these tend to be the most scalable. For this reason we focus on the OS variant of CBMs, where cell shape is not explicitly resolved but cells are instead modeled as spheres which can potentially overlap. Figure 1 illustrates the main idea. As depicted, cell connectivity is defined using a simple maximum interaction radius.

**Figure 1:**
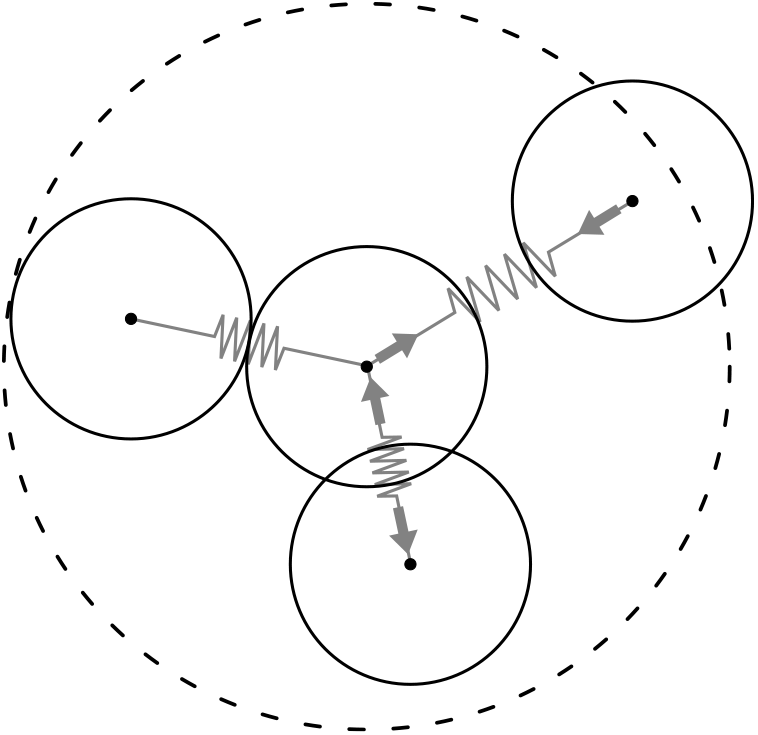
Illustration of the main idea of centre-based models, in particular the OS variant. Cell midpoint positions are tracked and the shape is assumed to be spherical, potentially overlapping with its neighbours. Cells move according to pairwise mechanical interactions, which can be repulsive (when overlapping) or adhesive (when non-overlapping but close). We assume that mechanical interactions are limited within a fixed distance from the cell centre, represented here by the dashed circle. All cells within this radius are assumed to interact with the cell in the centre.

Although several of the available frameworks for simulating CBMs are open and extensible, their large feature set, their focus on supporting modelers, and corresponding code complexity make them less suitable for rapid experimentation with numerical parameters. Here we study relatively simple models of cell mechanics with the need for full control of all the numerical settings. Therefore, we developed our own centre-based code in Python to ensure consistency across the numerical experiments. To facilitate reproducibility and in the hope that it will be useful for others for similar studies, the software is publicly available at: https://github.com/somathias/cbmos.git. The following subsections detail the implementation assumptions we consider in this study.

### 2.1 Update equation for the cell positions

Let **x**_*i*_ denote the midpoint coordinates of the *i*th cell and *m_i_* its mass. Applying Newton’s second function to the cell movement we obtain

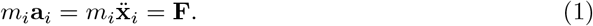

We assume that two types of forces act on the cell. First, a drag force **F**_drag_ stemming from the cell’s friction with the viscous environment is directed against the cell’s direction of motion. This force is proportional to the cell’s velocity,

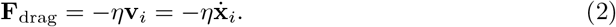

Secondly, neighbouring cells exert mechanical forces on the cell. In the simplest case these involve repulsive forces due to limited cell compressibility, but they usually also include cell-cell adhesion. Interactions are assumed to be pairwise and symmetric, i.e.

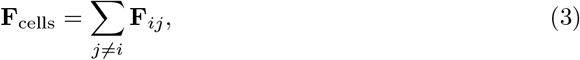

where the sum runs over all neighbours, excluding the cell itself. Which cells are considered neighbours depends on the definition of cell-cell connectivity, see Section 2.3 below for details. The resulting governing equation for the cell midpoint motion reads

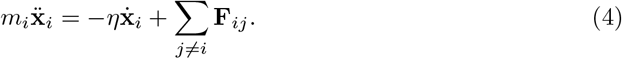

This is a system of second-order ordinary differential equations governing the cell centre positions, with one equation for each degree of freedom. As the microenvironment for the cells has a very low Reynolds number (the Reynolds number describes the ratio of inertial to viscous forces) [39], inertial effects such as acceleration are neglected. This commonly applied “inertialess” assumption, 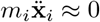, reduces the update equation to the system,

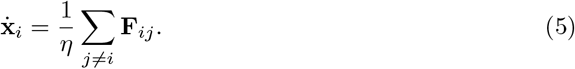

For increasing levels of physical realism in the simulations, Equation (5) might be complemented with additional terms representing friction between cells, repulsive and adhesive interaction between the cell and the extracellular matrix and cell migratory behaviour [9, 7]. The force-based formulation considered here has become the standard way of implementing CBMs and is favored over energy-based formulations (numerically studied using Monte Carlo methods such as the Metropolis algorithm [40, 35]) due to the straightforward interpretation of the time scale and a more intuitive way of treating custom cell interactions [7].

### 2.2 Numerical methods for solving the update equation

For an initial value problem stated as

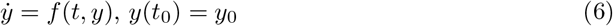

a numerical scheme provides an approximation for function values *y_n_* ≈ *y*(*t_n_*) at discrete time points *t_n_*, *n* = 1, …, *N*. The simplest numerical scheme, the forward Euler method, calculates the next function value by taking a step in the direction of the current gradient [41],

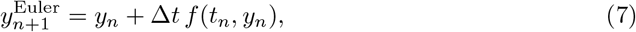

where Δ*t* is the step size. If the step size is chosen constant over the time interval *t*_0_ to *t_N_*, then Δ*t* = (*t_N_* − *t*_0_)*/N*. The forward Euler method is a first-order scheme, meaning that as long as Δ*t* is sufficiently small, the local error in one single time step is proportional to Δ*t*^2^ and the global approximation error at *t_N_* thus proportional to Δ*t*. Roughly speaking, halving the step size for a first-order scheme makes the solution twice as accurate.

Higher order schemes improve the convergence rate at the cost of additional function evaluations. An example is the midpoint rule [42],

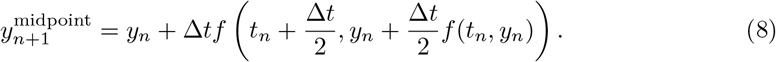

The midpoint rule is a second-order method, which means that halving the step size divides the error by four. Hence, less steps are needed to achieve a given accuracy compared to the forward Euler method, although each single step will be more costly.

Both methods are one-step methods, meaning that they calculate the function value at the next time point based on only the current function value. Multi-step methods additionally take past function values into account. One of the simplest two-step methods is the Adams-Bashforth method [41],

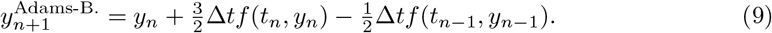

In addition to the order of the scheme, the schemes’ stability is an important characteristic. A numerically unstable solution will oscillate and grow without bounds, even though the true solution does not. In this case the step size needs to be reduced to recover a stable, bounded solution. The stability region depends on both the numerical scheme and on the ODE problem to be solved and this imposes an upper bound on Δ*t* for which the scheme can be used. The forward Euler method has very poor stability properties and may hence require very small time steps.

### 2.3 Definition of cell connectivity

The right-hand side in Equation (5) involves the evaluation of pairwise interaction forces for all cells *j* = 1, … , *M* in a neighbourhood of the cell with midpoint **x**_*i*_. In the overlapping spheres variant of CBMs that we consider, cell connectivity, i.e. which cells are allowed to mechanically interact, is defined via a maximum interaction distance (see Figure 1). This can be efficiently implemented in two and three dimensions. For a large number of cells the naive implementation can be improved by introducing so-called bounding boxes that keep track of closely located cells so that not all pairwise distances need to be evaluated in each step [43]. It is well-known that defining cell connectivity using only distance information between cells can lead to the problem of collapsing volumes [38]. In a densely packed population, newly divided cells can potentially interact with their next-to-nearest neighbours, resulting in the adhesive forces exerted by their (many) next-to-nearest neighbours becoming larger than the repulsive forces exerted by their (few) direct neighbours and thus compressing the overall volume of the population. Different force functions show a varying sensitivity to this issue, the linear spring being particularly problematic [38].

It is also possible to take topological information into account to ensure that cells only interact with neighbours with which they are in direct contact, i.e. that no other cell is located between them. This is done in the Voronoi variant of the CBMs using a Voronoi tesselation of the cell midpoints where cells are considered in mechanical contact if they share an edge in the associated Delaunay triangulation [1, 44, 45, 46]. The Voronoi polygons can then also be used to estimate cell shape and volume. This approach makes the Voronoi variant-CBM more robust to the issue of collapsing volumes at an increased computational cost compared to the OS variant [21, 38].

### 2.4 Time-driven vs. event-driven simulation

In addition to cell mechanics, a CBM simulation usually consists of a number of cell events, such as cell division, cell death, and potentially subcellular events such as transition in cell cycle models. Typically, simulations proceed through a split-step scheme with fixed time steps for each main model feature (mechanics, molecular diffusion, cell events etc.)[27]. This time-driven approach is a good way of handling systems with different physics and with different timescales, but it also introduces an additional level of approximation, a splitting error. This error in general is hard to estimate.

As an alternative, event-driven simulation methods maintain a list of time points at which events affecting the system will take place [47]. In the context of the simulation of cellular systems, this allows to simulate the mechanics until the next event time, apply the corresponding cellular event to the cell population and then continue with simulating the mechanics. This is possible as long as events can be scheduled in advance, e.g. by drawing a division time from a suitable distribution at cell birth, and do not depend on the current cell or population state. We chose this latter approach since it accurately resolves the cellular dynamics of the population for our model problems.

### 2.5 Cell cycle model and cell division implementation

In general, CBMs employ cell-cycle models of varying complexity to model the proliferation of cells, ranging from choosing a uniformly distributed cell cycle duration to nutrient- or contact-based cell cycle models [1, 9, 29, 4]. Furthermore, there exist different algorithms of implementing the cell division process itself. Biologically, this process involves the formation of a contractile actin ring at the division plane of the mother cell. As this ring contracts, the cell is gradually deformed leading to the formation of two daughter cells that eventually pinch off into two adhering cells of roughly half the volume of the mother cell. Since cell shapes are not explicitly modeled in the CBM, the details of this process cannot be captured directly. Instead, different types of approximations are employed. A thorough discussion on the various aspects of cell division algorithms, including a summary of additional commonly modeled mechanisms in the cell cycle such as contact inhibition and apoptosis can be found in [23]. We here briefly discuss two approaches.

In [9], two main phases of division are recognized. A cell entering the cell cycle first shows spherical growth. Then, when it splits into two daughter cells, they are treated as a dumbell in which the cells gradually separate by increasing the distance to each other. Alternatively, cells are treated as a dumbell directly when they enter the cell cycle and spherical growth and separation happens simultaneously [37]. Dumbell formation, although somewhat close to the biological processes in concept, introduces additional complexity in the model due to breaking of spherical symmetry [9].

A simpler and more commonly used algorithm is to instantaneously place two newly formed daughter cells of a smaller radius close to each other (with an overlap) in the space previously filled by the mother cell [44]. The cells will then relax into mechanical equilibrium with their surroundings according to the specific force function dynamics used in the simulation, as well as increase their radius to the size of the former mother cell. This, or close variants of it, is also the algorithm used in several recent high-performance and parallel implementations [48, 27, 49]. Implementation-wise this is a very simple and efficient algorithm. However, it may lead to locally and transiently high, unphysical force values right after cell division [44, 23]. This can be seen as a consequence of the fundamental model assumption underlying the CBM, i.e. the assumption of pair-wise cell interactions not being valid in highly compressed scenarios [23]. In [50] a multiscale approach based on deformable cell model simulations is used to correct forces for highly compressed structures. We will instead investigate how the population level behaviour is affected if these high local forces are not correctly resolved numerically.

For our numerical study, we consider only cell division events and do not model the cell cycle explicitly. We adopt the simplified division algorithm detailed above and in [44], but without accounting for cell growth. We let cells divide instantaneously into two overlapping daughter cells of volume equal to the mother cell, see Figure 2. Unless stated otherwise, cells divide after a random time *τ* drawn from a normal distribution with a mean of twenty-four hours and a standard deviation of 1 hour. The cell division direction is chosen randomly according to a uniform distribution of the angle.

**Figure 2:**
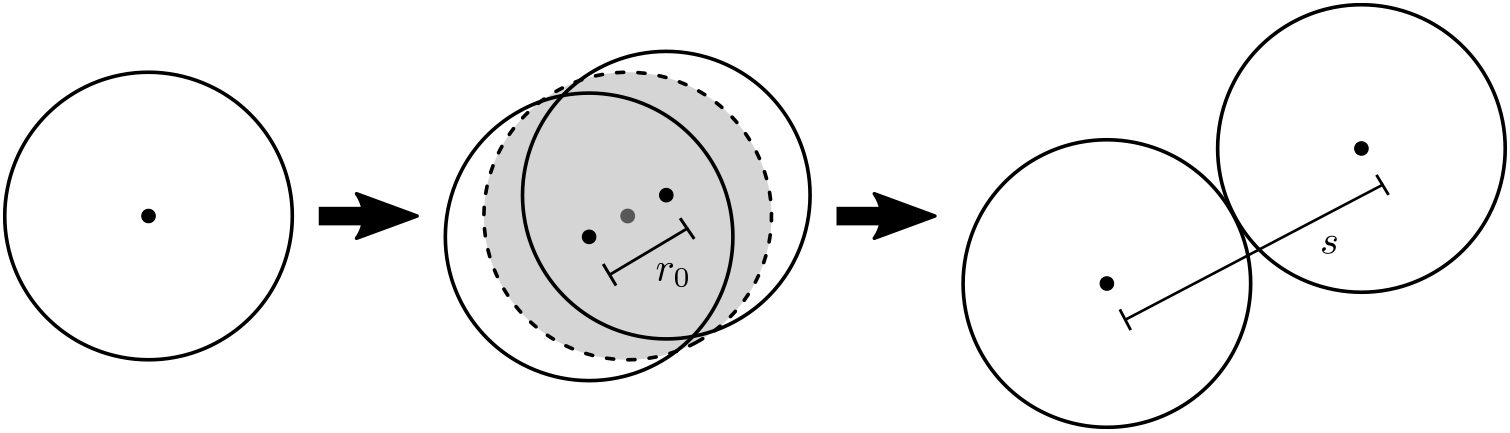
Illustration of the cell division steps in our implementation. Left: A cell ready to proliferate. Middle: Placement of two daughter cells of radius equal to that of the mother cell. The former position of the mother cell is the shaded region. The distance between the centres of the daughter cells is the initial separation *r*_0_. The cell division direction is chosen randomly. Right: After mechanical relaxation cell midpoints are separated by the rest length *s*.

## 3 Cell-cell interaction forces

The cell-cell interaction force **F**_*ij*_ in Equation (5) describes how cells interact mechanically and thus determines the biophysical behaviour the model can capture. In this study we consider force functions based on the following assumptions:

1. The pairwise force can be written as 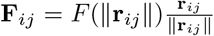, where **r**_*ij*_ = **x**_*j*_ − **x**_*i*_ and *F* is a scalar force. This means the force acts only in direction of the vector between cell centres and its magnitude depends only on their distance.
2. There exists a distance between cell midpoints (the rest length *s*) at which cells are in equilibrium, i.e. they exert no forces on each other and hence *F* (*s*) = 0.
3. Cells have limited compressibility and push each other for centre-centre distances smaller than the rest length to minimize their overlap, i.e. *F* (*r*) < 0 for *r* < *s*.
4. Forces are either zero or cells exert a pull on each other due to cell-cell adhesion forces for distances larger than the rest length, i.e. *F* (*r*) ≥ 0 for *r* > *s*.
5. There exists a maximum interaction range *r_A_* beyond which cells do not interact mechanically with one another, i.e. *F* (*r*) = 0 for *r* ≥ *r_A_*.

The simplest force function fulfilling these assumptions is a linear spring, used in early works by Drasdo and others [51, 52]. For the linear spring, the force is assumed to be directly proportional to the distance between cell midpoints, vanishing at the rest length. While this behaviour may be reasonable for small distances, it results in unphysically strong long-range interactions if extended to distances larger than the rest length. At the same time, a linear force does not result in very large repulsive forces when cells are very close, i.e. cells are highly compressible. This can lead to the problem of collapsing volumes, where a cell population collapses on itself under strong compression [38]. Additionally, the linear force will feature a large discontinuity at the maximum interaction range which can lead to further numerical difficulties. For both these reasons, force functions are favoured which feature (i) stronger repulsive interactions for cell separation distances close to zero and (ii) adhesive interactions that vanish for large cell distances. Examples of force functions with these properties include classical potentials such as the Morse or the Lennard-Jones potential [53], physically motivated force functions such as Hertz [54], and a number of mathematical functions built by extending the linear force with different terms. Table 1 summarizes the mathematical equations for several cell-cell interaction functions used for CBMs and examples of their references in literature.

**Table 1:**
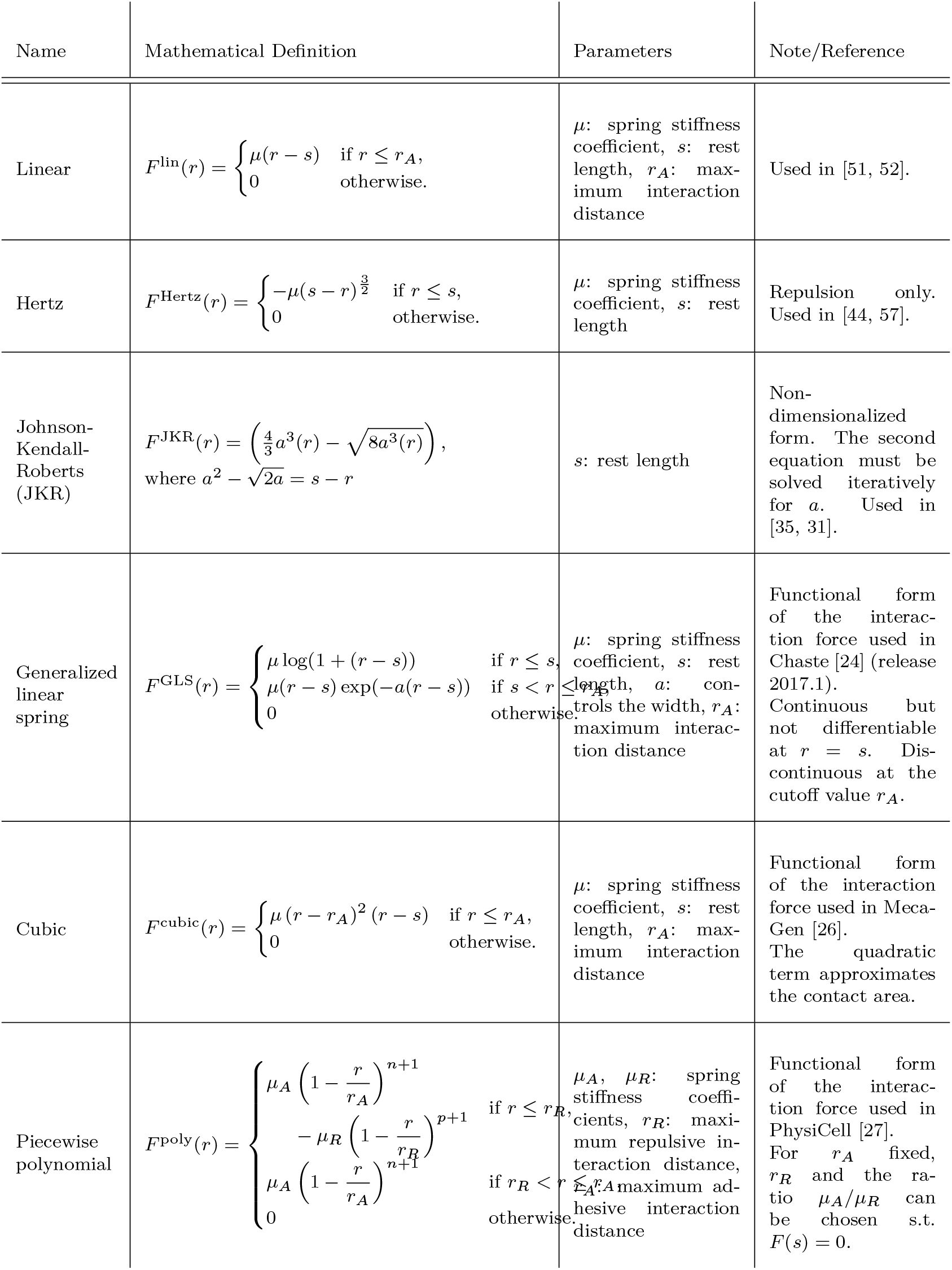
Overview over cell-cell interaction functions used for CBMs. These scalar functions are extended to 2D or 3D by multiplication with the normalized direction vector between cell midpoints, i.e. we define the force vector as 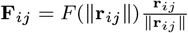, where **r**_*ij*_ = **x**_*j*_ − **x**_*i*_. In this paper, we focus on comparing the GLS, cubic and piecewise polynomial forces.

Note that we do not consider a hysteresis effect where the adhesion between cells depends on whether they were in contact before or not. Such an effect is modeled by the Johnson-Kendall-Roberts (JKR) model for the deformation of elastic bodies, which has been confirmed experimentally to be valid for biological cells under certain conditions [55]. Unfortunately, the JKR force is computationally very expensive since it requires the solution of an implicit equation in order to recover the force for a given centre-centre distance [35]. To decrease the computational requirements, the JKR can be approximated by a third order polynomial as done in CellSys [31, 56]. It would generally be possible to include a kind of hysteresis effect for the force functions we consider here, by applying an adhesive force only if cells are moving apart, but not when they are approaching each other. We leave this approach and its evaluation for future work.

In what follows we focus on the cubic, the piecewise polynomial and the generalized linear spring force. These pair-wise interaction forces are those implemented as default forces in the open-source software packages MecaGen [26], PhysiCell [27] and Chaste [24] and therefore it is likely that these functions are those that modellers new to the field come into contact with first. In the following subsections we describe the three chosen force functions and study if and how they can be parameterized based on pairwise interactions in order to produce similar biological behaviour at the population level. We will then use this parameterization as a starting point for a quantitative comparison of key numerical properties of CBM simulations using these force functions in Section 4.

### 3.1 Description of the mathematical form of the force functions and their parameters

In their respective software packages, these force functions are stated and parameterized very differently, so we start by describing each of them within a consistent notation and parameterization. In what follows, let *r* denote the centre-centre separation distance between a pair of cells, *s* the rest length and *r_A_* the maximum (adhesive) interaction distance. The maximum interaction distance *r_A_* and the rest length *s* are parameters that relate to the biophysical properties of cells and are treated as fixed parameters given as initial data to a simulation.

The first of the three force functions we have chosen, the cubic force, combines a linear spring term with a quadratic term approximating the contact area of two overlapping spheres. It is used in the software MecaGen [26] and can be written as

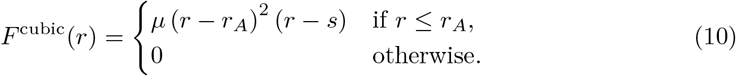

Here, the only free parameter *μ* denotes the spring stiffness parameter. In order to ensure differentiability of the force at the rest length, i.e. for *r* = *s*, *μ* has to be independent of whether cells are overlapping (*r* < *s*) or not (*s* ≤ *r* < *r_A_*). Larger values of *μ* increase the steepness of the repulsive interaction part, making cells behave more rigidly. At the same time, the magnitude of the adhesive interaction is increased, see Figure 3a.

**Figure 3:**
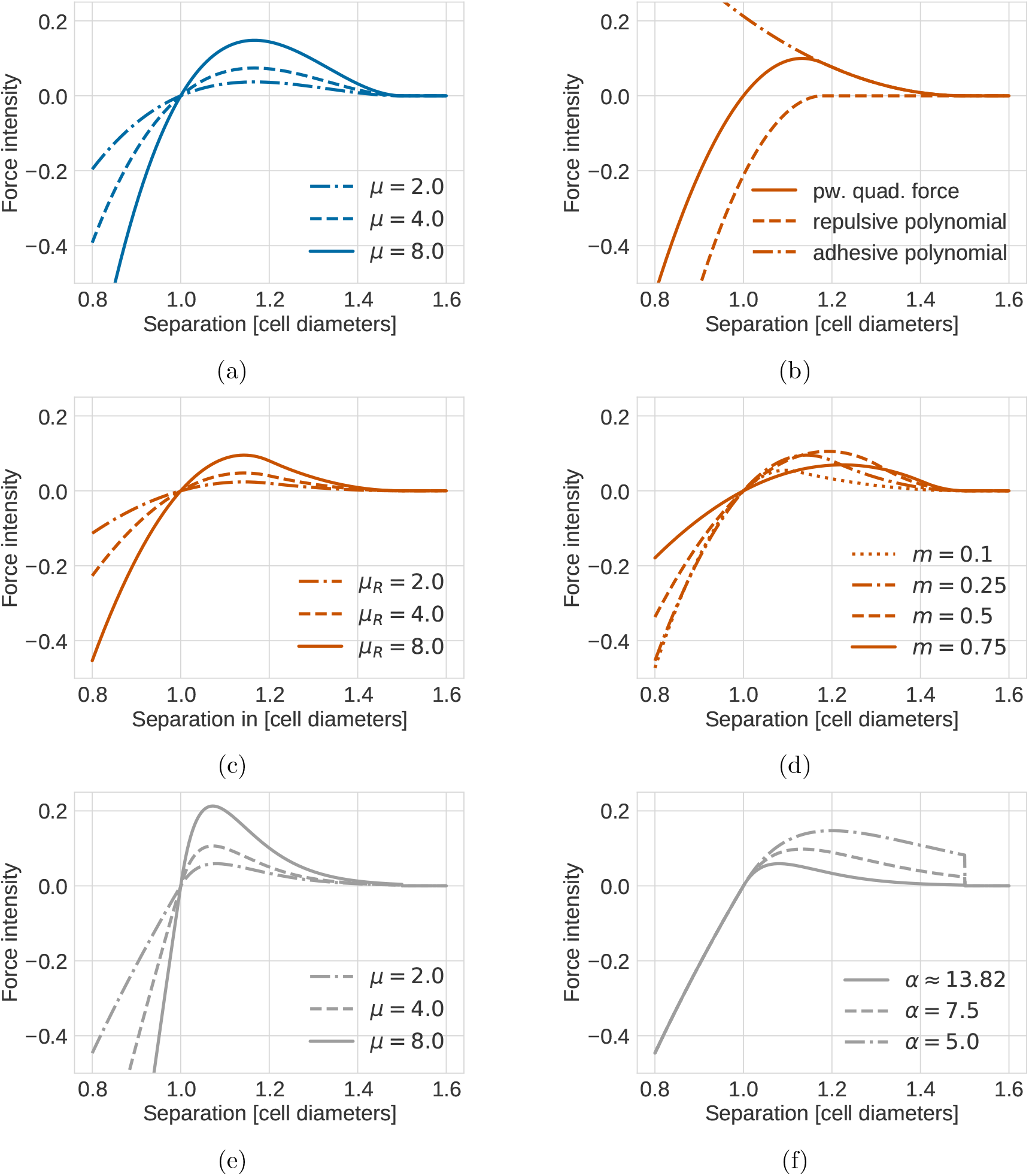
Intensity of the different forces as a function of cell separation distance. (a) Intensity of the cubic force shown for different spring stiffness values *μ*. (b–d) Intensity of the piecewise quadratic force. (b) The repulsive and adhesive polynomials are shown individually. Parameters were chosen as *μ_R_* = 9.1, *m* = 0.21. (c) A fixed ratio *m* = 0.21 was chosen and different spring stiffness values *μ_R_*. (d) A fixed spring stiffness value *μ_R_* = 8.0 was chosen and different ratios *m*. In contrast to the cubic and GLS force, the PWQ force allows for varying the repulsive and adhesive spring stiffness parameter independently within a certain range. (e–f) Intensity of the GLS force with (e) different spring stiffness values *μ* and (f) *μ* = 2.0 and different values of the bredth *α*. Here, for the largest *α*, as well as in (e) *α* is chosen according to Equation (14).

The second force function, the piecewise polynomial force is constructed as the sum of a positive adhesive and a negative repulsive polynomial, the support of which overlap, see Figure 3b. As a default quadratic polynomials are used in PhysiCell [27]. We take the same approach, leading to the formula,

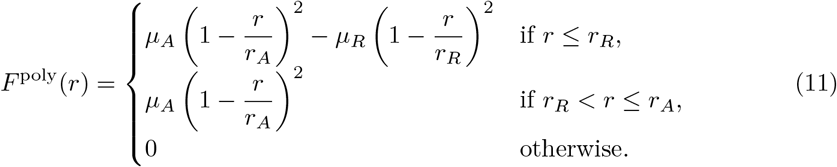

Here, *r_R_* denotes the maximum repulsive interaction distance, and *μ_A_* and *μ_R_* are the adhesive and repulsive spring stiffness parameters. Note that in general *s* < *r_R_* < *r_A_* and *μ_A_* < *μ_R_*. As stated earlier, we require that *F* (*s*) = 0, i.e. that the force vanishes at the rest length. This effectively fixes the maximum repulsive interaction distance *r_R_* as a function of the ratio 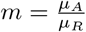 of the adhesive to the repulsive spring stiffness,

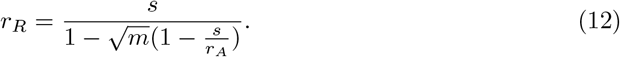

Consequently, the piecewise quadratic function has two free parameters, *μ_R_* and *m*. Figures 3c and 3d show the force intensity as a function of cell separation distance for different parameter combinations. We observe that for a fixed ratio *m*, increasing the repulsive spring stiffness *μ_R_* increases the magnitude of the force, similarly to the cubic force function (see Figure 3c). On the other hand, Figure 3d shows that the piecewise polynomial force function lets the modeler tune the level of adhesive interaction for a fixed repulsive strength within a certain range, which is not possible for neither the cubic nor the generalized linear spring.

Last but not least, the generalized linear spring (GLS) force as implemented in Chaste [24] uses a logarithmic force for the repulsive regime. For cell distances larger than the rest length it extends this force with a linear spring term multiplied with an exponential term to ensure decay of the force for large distances.

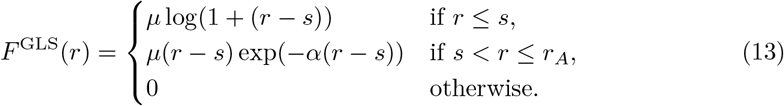

Contrary to the cubic and the piecewise quadratic force function, the GLS force is not constructed in a way that it vanishes for cell separation values larger than the maximum interaction distance. Instead the force is just cut off at that value, leading to a discontinuity at *r* = *r_A_*.

The GLS force has two free parameters, the spring stiffness *μ* and the parameter *α* which controls the width of the exponential decay in the adhesive regime. Similarly to the cubic function, ensuring differentiability forces the spring stiffness *μ* to be the same for the repulsive and the adhesive regime. Figures 3e and 3f show the intensity of the GLS force for different values of *μ* and *α*. We observe that as with the other force functions, increasing the spring stiffness *μ* leads to stronger repulsive and adhesive interactions (for a fixed value of *α*) (see Figure 3e). Decreasing *α* for a fixed spring stiffness value *μ* increases the amplitude of the adhesive interaction while not affecting the repulsive interaction (see Figure 3f). However, it leads to a larger discontinuity at the maximum interaction distance *r_A_* which will affect the numerical properties of the GLS force as we will see in Section 4.3. We will choose *α* dependent on *μ* in a way to ensure that the magnitude of the force is small at the maximum interaction distance *r_A_*, requiring that *F* (*r_A_*) ≤ *ϵ* = 10^−3^. Then we obtain

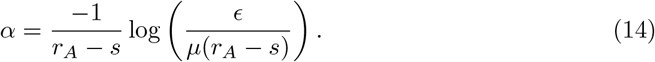

Unless stated otherwise, we consider the spring stiffness *μ* as the single free parameter of the GLS force.

Having described the force functions individually, we will now move on to compare their qualitative behaviour.

### 3.2 Comparison of the qualitative behaviour for a fixed relaxation time

The three studied force functions all have different functional forms and a varying number of free parameters. A natural question faced by a modeler is to what extent the force functions are interchangeable, i.e. whether they will result in simulations leading to the same biological conclusions given an appropriate parameterization. To study this, we first use a simple model system consisting of the fundamental unit of two interacting cells. We then check that slight differences at the level of pairwise interaction do not have a strong impact on a population level metric for two dimensional planar growth.

In all experiments, we (i) let cells have a fixed radius of *R* = 0.5 cell diameters and (ii) fixed the rest length *s* and the maximum interaction distance *r_A_* to the same value for all force functions. We chose the rest length to equal one cell diameter, i.e. *s* = 2*R* = 1.0 cell diameter. To ensure that a cell configuration placed at Cartesian coordinates relaxes to a honeycomb-like configuration, but that next-to-nearest neighbours do not interact at rest, the maximum interaction radius has to be chosen between 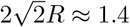 and 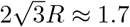 cell diameters, as illustrated in Figure 4. We used *r_A_* = *s* + *R* = 1.5 cell diameters.

**Figure 4:**
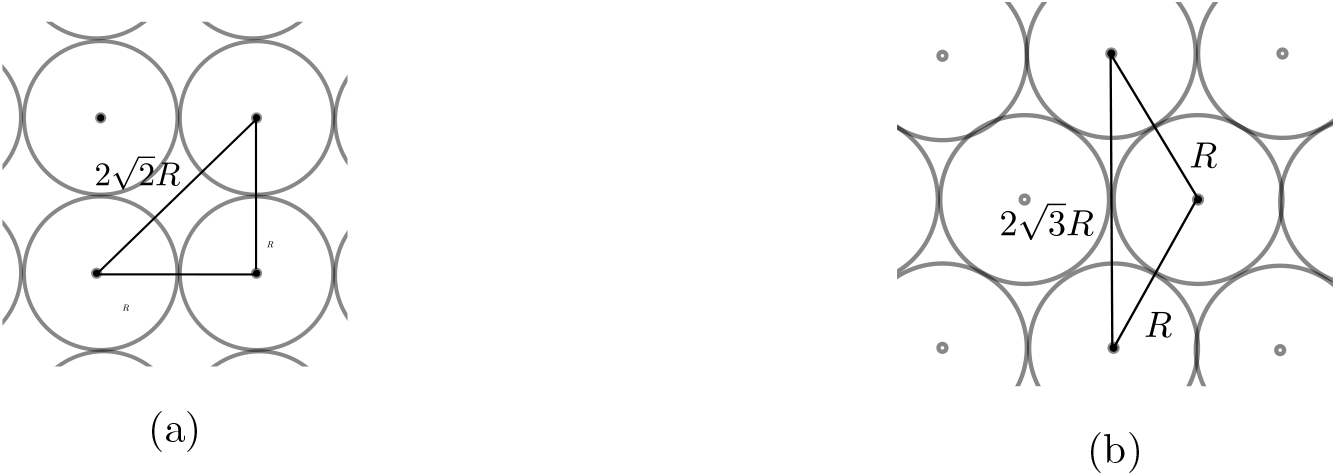
Illustration of cell centre-centre distances for populations arranged on (a) a cartesian grid and (b) a hexagonal lattice in a so-called honeycomb configuration.

To select the remaining free parameters we assumed that all force functions should result in the same pairwise relaxation time after cell division. This can be seen as a basic requirement of interchangeable force functions for a proliferating system. However, it leads to differences in the transient relaxation dynamics in the repulsive regime, and also to differences in force magnitudes in the adhesive regime. First, we illustrate these differences. Then we study their impact on a population level metric for a system of proliferating cells, the population radius, and examine whether the fit can be improved by changes in the parameterization.

#### 3.2.1 Pairwise relaxation dynamics

First, we considered two overlapping cells as the relax to mechanical equilibrium right after cell division. The cells were placed at a fixed separation distance *r*_0_ = 0.3 cell diameters and the system was evolved until the overlap was eliminated due to the repulsive forces (see Figure 2 for an illustration). Parameters were chosen to ensure that for all force functions the relaxation time *t*_0_ is equal to one hour. Here, we defined the relaxation time as the time such that the distance between cells is equal to 99% of the rest length *s*. The parameter values were determined numerically using the miminize function of the scipy.optimize library in combination with our CBMOS simulation code to calculate the separation distance as a function of the spring stiffness *μ*. The resulting parameter values are stated in Table 2.

**Table 2:** Parameter values for a fixed relaxation time *t*_0_ = 1 *h* after cell division. Free parameters are printed in bold. The spring stiffness values *μ_cubic_*, *μ_R_* and *μ_GLS_* have been determined numerically by minimizing the difference between the separation distance *r* and 99% of the rest length *s* at *t*_0_, i.e. min_*μ*_ ‖*r*(*t*_0_; *μ*) − 0.99*s*‖. Here, the minimization was done using the BFGS algorithm implemented in the minimize function of the *scipy.optimize* library [58, 59], where the separation distance was evaluated for different spring stiffness values *μ* using our CBMOS simulation code. Values were rounded to two decimal values. For the piecewise quadratic force, the minimization was done jointly over *μ_R_* and the ratio *m*. The adhesive spring stiffness was then chosen as *μ_A_* = *m* × *μ_R_*. The values for *r_R_* and *α* where chosen according to Equations (12) and (14).

| Parameter | Description | Value |
| --- | --- | --- |
| **$\mu_{\text{cubic}}$** | spring stiffness, cubic force | 5.7 |
| **$\mu_R$** | repulsive spring stiffness, pw. quad. force | 9.1 |
| **$m$** | ratio of adhesive to repulsive spring stiffness, pw. quad. force | 0.21 |
| $\mu_A$ | adhesive spring stiffness, pw. quad. force | 1.91 |
| $r_R$ | repulsive interaction distance, pw. quad. force | 1.18 |
| **$\mu_{\text{GLS}}$** | spring stiffness, GLS force | 1.95 |
| $\alpha$ | breadth of exponential, GLS force | 13.76 |

Figure 5a shows the separation in cell diameters as a function of time for the different force functions. Over time the separation between the cells correctly approaches the rest length for all force functions. The rate at which the overlap is eliminated, however, differs between force functions. It decreases fastest for the cubic function and slowest for the GLS force. This means that cells behave more or less rigidly depending on the force function, even if they eliminate their overlap completely within the same time duration (same relaxation time). We will see in Section 4.1 that these differences in the stiffness of the system are reflected in the numerical stability bounds.

**Figure 5:**
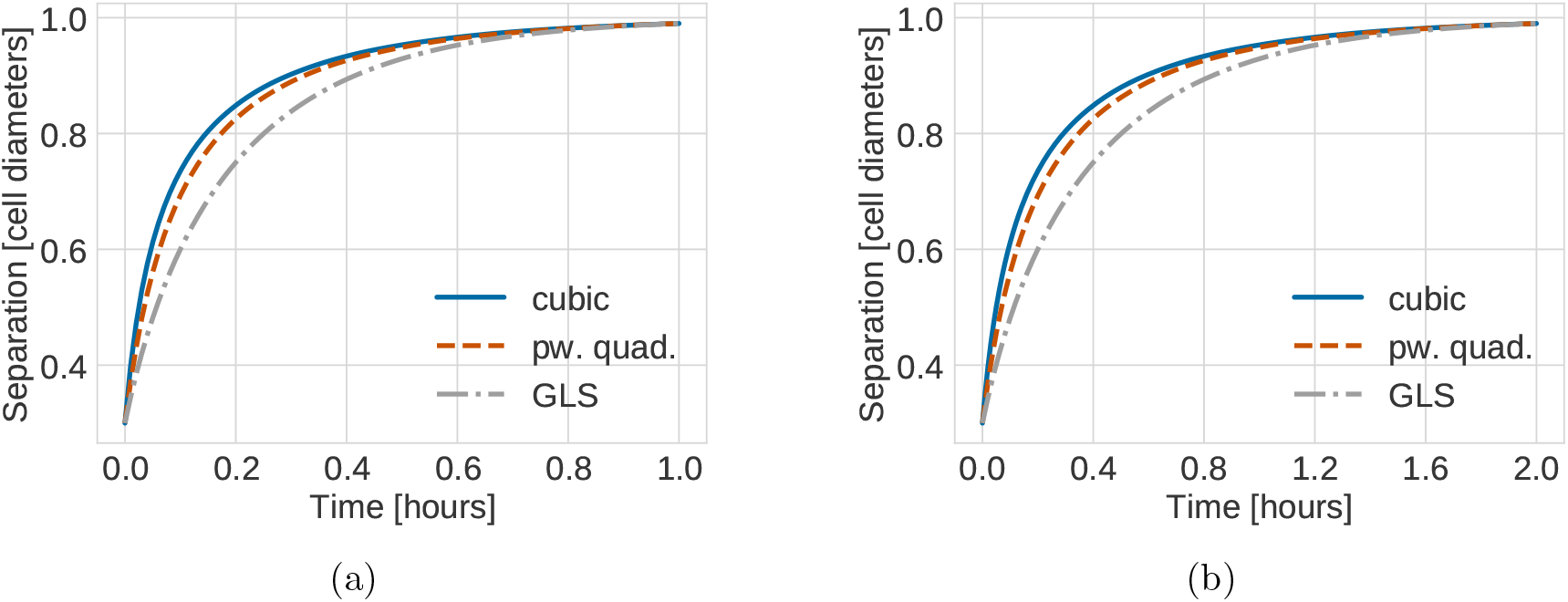
Relaxation dynamics between two cells initially placed at a fixed separation of 0.3 cell diameters. The parameters have been chosen such that the separation equals 0.99 cell diameters after (a) 1 hour and (b) 2 hours.

Doubling the relaxation time by halving the spring stiffness *μ* does not change the qualitative dynamics but scales the time scale, compare Figure 5b to 5a. This can be seen analytically from stating the simple ODE for the distance between a pair of mechanically relaxing cells after division,

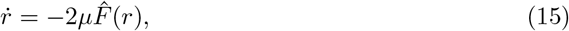

where we have made the dependence on the spring stiffness *μ* explicit. Rescaling time as *τ* = *μt* leads to 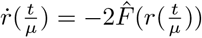, meaning that increasing *μ* speeds up the dynamics and decreases the relaxation time. Consequently, a longer relaxation time decreases the stiffness of the system at the cost of having to run simulations for a longer time. In the following we considered the time scale on which mechanical relaxation happens to be (arbitrarily) fixed as 1 hour.

Next, we asked how cells interact in the adhesive regime when parameterized to have the same fixed relaxation time. Figure 6a shows the intensity of the force functions as a function of cell separation. We observe that the cubic and the piecewise quadratic force agree well in the repulsive regime, i.e. for *r* < *s*. However, the GLS force does not agree well for separation values of *r* < 0.9 cell diameters. It also has a much lower maximum amplitude in the adhesive regime, i.e. *r* > *s*, and for smaller separation values than the cubic and the piecewise quadratic force. While the magnitude for the latter two forces is similar, the piecewise quadratic force shows a steeper decrease to zero magnitude at the maximum interaction distance. Figure 6b shows the resulting pairwise dynamics for cells located at different centre-centre distances larger than the rest length. We observe that the transient dynamics differ qualitatively. In the next section, we will turn to investigate how this affects a freely growing monolayer under compression from high proliferation, where the repulsive interactions play the major role, but adhesion between next-to-nearest neighbours is also present.

**Figure 6:**
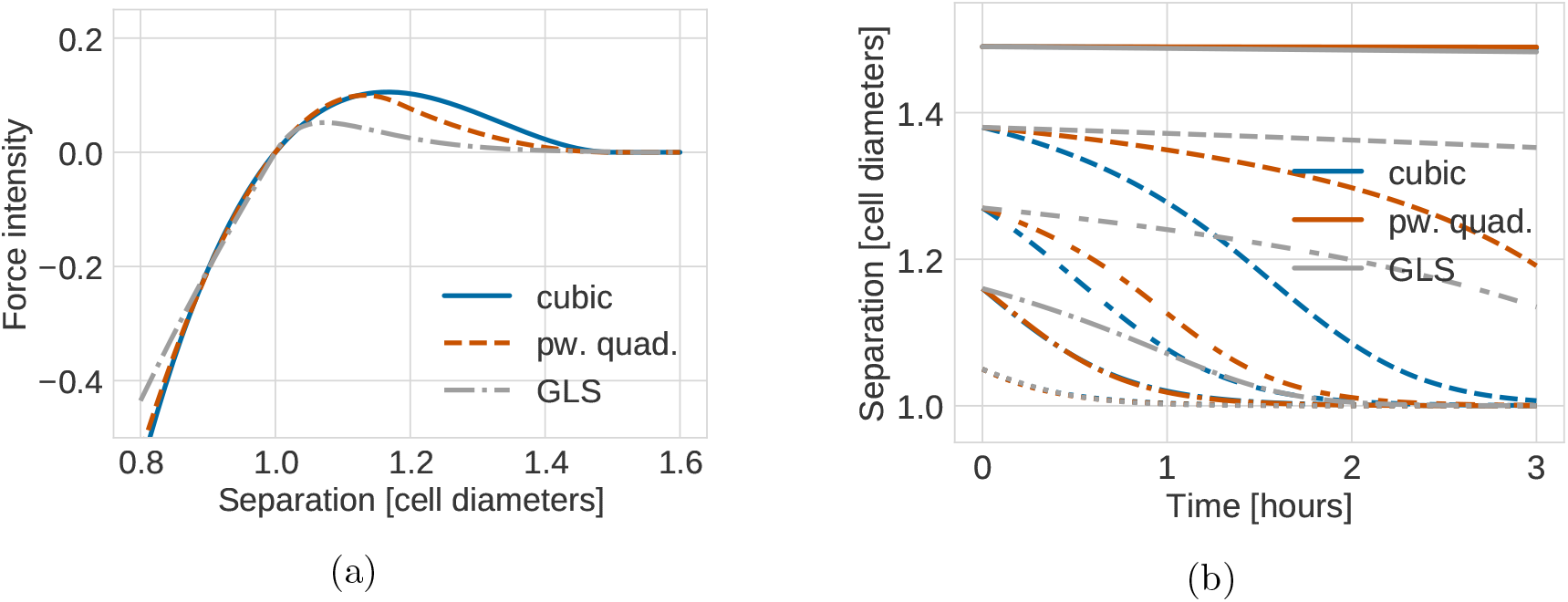
(a): Intensity of the different force functions as a function of cell-cell separation for the adhesive regime. (b): Relaxation dynamics between adhering cells under the different force functions. Two cells are initially placed at different distances within the maximum interaction distance of 1.5 cell diameters. Different linestyles correspond to different initial separation distances between the two cells. Parameters for both have been chosen such that the force functions have the same relaxation time after cell division as shown in Table 2.

#### 3.2.2 Impact of the parameterization on the radius of a monolayer population relaxing after intense proliferation

As discussed in the previous subsection, calibrating the force functions to agree on the mechanical relaxation time is a simple and computationally cheap procedure. But since the parameterization is based on a summary statistic, the forces disagree quantitatively both in the strongly compressed repulsive regime and in the adhesive regime. The timescale for which the forces show large disagreement in the repulsive regime is relatively short compared to the relaxation time. As can be seen in Figure 6a the force functions agree well in the repulsive regime at a separation of 0.85 cell diameters or more. In Figure 5 we see that this separation is reached in less than the first 30% of the total relaxation time for both the shorter and longer relaxation times. The disagreement in the adhesive regime is larger. In this section, we investigate how these disagreements affect the relaxation of a small growing monolayer. As we are interested in how the force functions compare when modelling mechanical relaxation due to proliferation of the tissue, we choose a setting of high compression by letting all cells divide simultaneously.

In Figure 8 we show force intensities for five parameterization strategies (left column) and corresponding simulation results for the population level radius of a growing monolayer in two dimensions (middle and right columns). In the middle column, we initialised 19 cells arranged in a honeycomb pattern. We then let all cells divide simultaneously at the beginning of the simulation and tracked the population radius over 10 hours (in-simulation time). During this time the resulting monolayer of 38 cells relaxed to a mechanical equilibrium configuration. Figure 7 shows an illustration of the procedure. For the right column, we did the same procedure for an initial honeycomb pattern of 37 cells, resulting in a final population size of 74 cells, approximately doubling the number of cells in our first experiment. This was done in order to empirically check the robustness of our results to the size of the growing monolayer. In both cases the population radius was calculated by taking the maximum distance of any cell to the centroid of the population and adding the radius *R* of a single cell.

**Figure 7:**
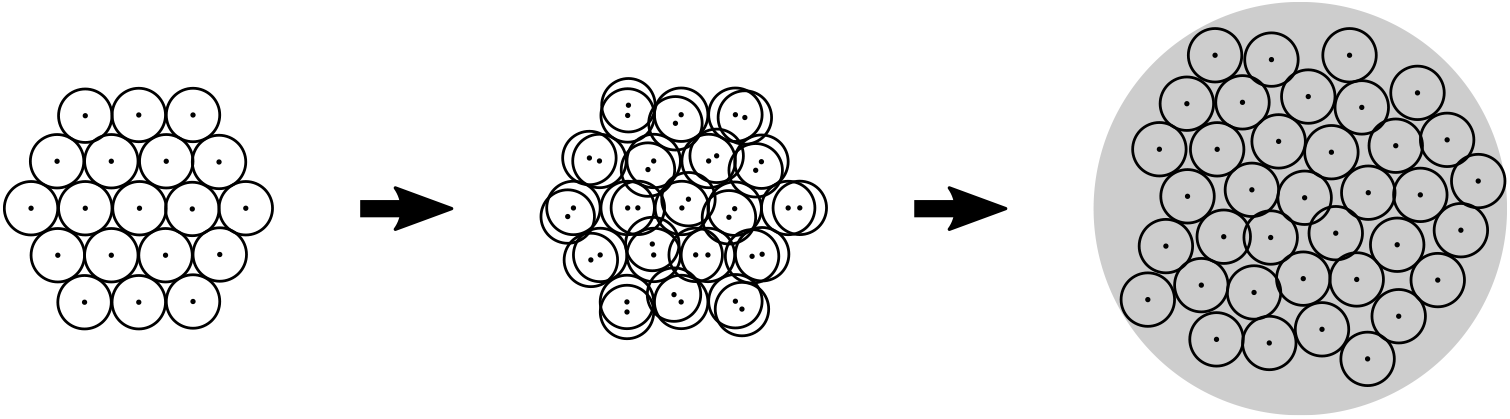
Illustration of a monolayer growing and relaxing under intense proliferation. Initially, 19 cells are placed in a honeycomb arrangement (left). Then all cells are simultaneously let to divide (middle). The resulting population of 38 cells is mechanically relaxed to equilibrium and the population radius tracked over the process (right). Note that due to adhesive forces present the equilibrium state can include slight overlaps between cells. The procedure is done similarly for an initial placement of 37 cells (not shown).

**Figure 8:**
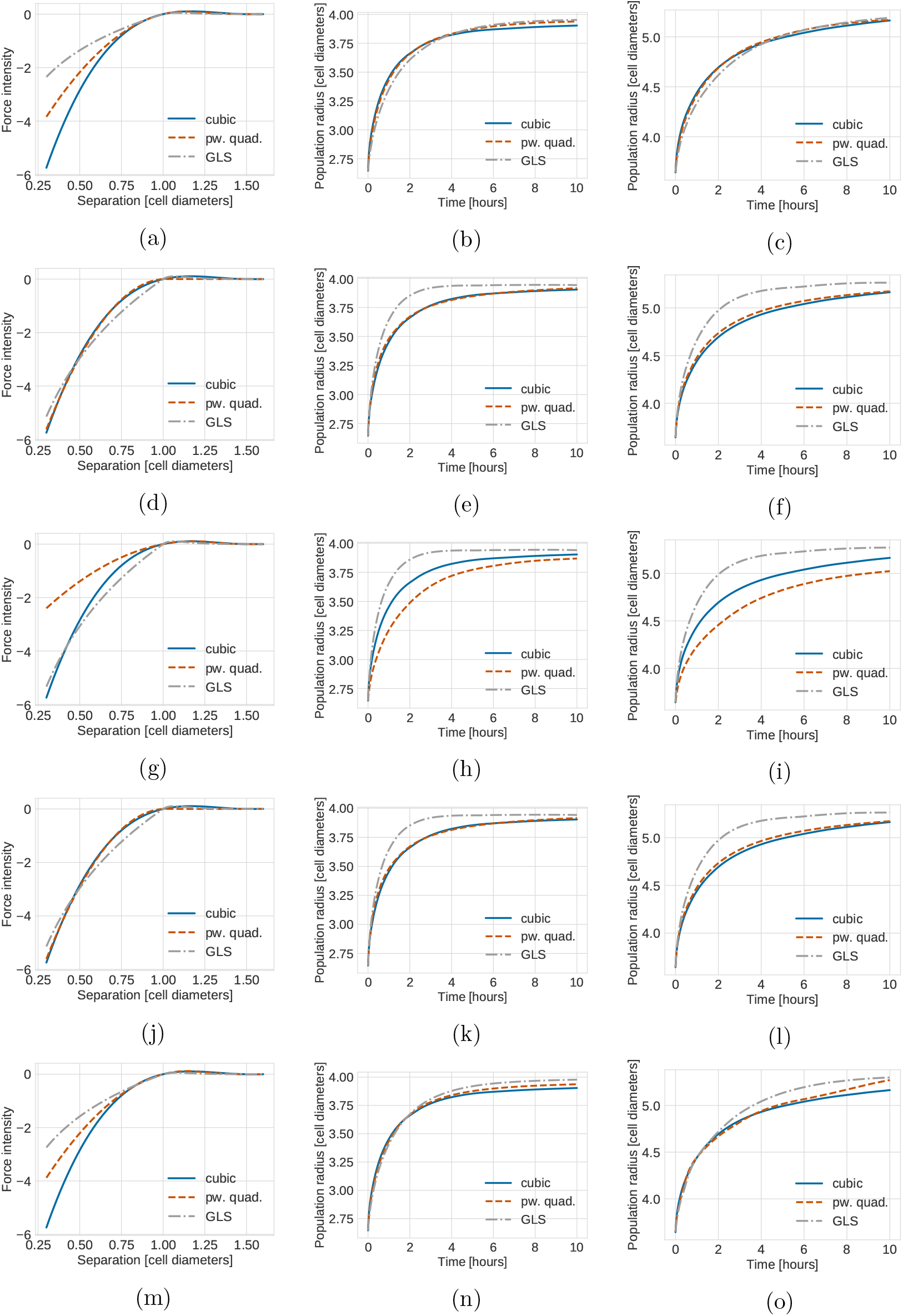
Plot showing force intensities for four parameterization strategies (left column) and corresponding simulation results for the population level radius of a growing monolayer in two dimensions of initially 19 cells (middle column) and of initially 37 cells (right column). The population radius for each time point was calculated as 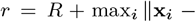 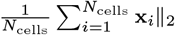 where *R* = 0.5 cell diameters is the radius of a single cell. The population radius was then averaged over 10 simulation runs with different random seeds. (a)-(c) Fit to relaxation time, (d)-(f) fit over repulsive range, (g)-(i) fit over adhesive range, (j)-(l) fit over complete range, (m)-(n) fit to population radius for 38 cells, (o) population radius for 76 cells using the parameter values fit to a population radius for 38 cells as shown in (m) and (n). The individual parameter values used can be found in Tables 2 and 3.

First of all, we note that our experiments showed that independently of the exact parameterization used, all three forces are robust to the issue of collapsing volumes discussed in [38], since in no case the population radius decreases to values smaller than that of the initial configuration (see all subplots of Figure 8).

Secondly, we observe that matching relaxation times results in good agreement for all force functions on the population level metric despite large force discrepancies in the highly compressed regime, Figures 8a, 8b and 8c. Empirically, this seems to hold true independent of the actual size of the monolayer, as the curves agree very well for both monolayer sizes of initially 19 and 37 cells. Fitting the force functions by minimizing the discrepancy over the entire repulsive regime works well for the cubic and the piecewise quadratic force but results in larger differences in population radius for the GLS force due to large force discrepancies in the medium-to-low compressed regime, Figures 8d and 8e. The corresponding strategy of fitting over the adhesive regime again results in worse agreement, Figures 8g and 8h. We also attempted to fit over both the repulsive and adhesive regime (Figures 8j and 8k) but saw no improvement compared to Figures 8d and 8e. This can be explained by the fact that in our experimental setup, where the monolayer relaxes after intense proliferation, most interactions will be of the repulsive kind.

Finally, we parameterized the functions by fitting to the actual population radius for the monolayer of initially 19 cells, Figures 8m and 8n. The strategy here was to numerically fit the population radius obtained from simulations using the piecewise quadratic and GLS forces to the population radius averaged over 10 simulations with the cubic force. Interestingly, this approach resulted in similar force parameters to those obtained by matching relaxation times, compare Table 2 and Table 3. The increase in computational time needed to do the same numerical optimization procedure for the experimental setup of initially 37 cells prohibited us from doing so. Instead we checked how the parameters from the smaller monolayer transferred to the larger monolayer in Figure 8o. In contrast to when fitting to the relaxation time, the population radius does not agree well between the different forces for this case.

**Table 3:** Parameter values for different fitting procedures. The spring stiffness values *μ_R_* and *μ_GLS_* were determined numerically by minimizing the difference between force function magnitudes for different ranges of cell separation values *r*. For the fit over the repulsive range *r* = [0.3, 1.0], for the fit over the adhesive range *r* = [1.0, 1.5] and for the fit over the complete range *r* = [0.3, 1.5]. In all cases, the cubic force with parameters as stated in Table 2 was used as a reference. The minimization was done using the BFGS algorithm implemented in the *minimize* function of the *scipy.optimize* library [58, 59]. Values were rounded to two decimal values, except for the ratio where they were rounded to three decimal values. For the piecewise quadratic force, the minimization was done jointly over *μ_R_* and the ratio *m*. The adhesive spring stiffness was then chosen as *μ_A_* = *m* × *μ_R_*. The values for *r_R_* and *α* where chosen according to Equations (12) and (14). Their values are stated here rounded to three decimals. For the fit to population radius the population dynamics were simulated for a given spring stiffness value using our CBMOS simulation code. Again, the dynamics obtained using the cubic force fitted to a relaxation time of one hour were used as reference.

| Fitting procedure | $\mu_R$ | $m$ | $\mu_A$ | $r_R$ | $\mu_{GLS}$ | $\alpha$ |
| --- | --- | --- | --- | --- | --- | --- |
| Fit over repulsive range | 11.35 | 0.002 | 0.023 | 1.015 | 4.27 | 15.332 |
| Fit over adhesive range | 8.13 | 0.462 | 3.740 | 1.292 | 4.43 | 15.406 |
| Fit over complete range | 11.33 | 0.002 | 0.023 | 1.015 | 4.27 | 15.332 |
| Fit to population radius of initially 19 cells | 9.76 | 0.260 | 2.538 | 1.205 | 2.28 | 14.078 |

In conclusion, for the initial stages of two dimensional monolayer growth, the three force functions we consider can all result in close macroscopic readouts (in this case the population radius) for translations of parameters such as repulsive spring stiffness. In this sense, they can be seen as interchangeable from a modelling perspective, at least for the model system we consider. Interestingly, fitting to the relaxation time leads to better agreement than when transferring parameters obtained from fitting directly to the population radius from a smaller to a larger population. Having established a way of effectively parametrizing the three forces using the simple pairwise metric of relaxation time, we now turn to discuss how the forces compare numerically. We will see that the fact that the force intensities differ in the highly compressed regime can have an impact on the numerical solution of the equations of motion.

## 4 Numerical properties of three popular force functions

Given a parameterization of the various force functions that result in similar population level behaviour, it is interesting to study the properties of the numerical solution of the corresponding ODE (6). In particular, we are interested in differences in the error as the system approaches equilibrium for given time step sizes Δ*t*, since this directly affects the simulation efficiency for any implementation of a CBM simulator. The large force gradients right after cell divisions (when cells might overlap to a large extent) causes the ODE system to be stiff. Commonly used explicit schemes such as the forward Euler method are simple to implement and avoid the expensive solution of large non-linear equations systems needed for implicit methods. However, explicit methods struggle with stiff systems because the time step will be forced to take small values in order to ensure stability and accurate resolution of the transient solution. We are primarily concerned with differences in the numerical properties depending on the force function chosen.

Similarly to the structure of Section 3 we start by examining the numerical stability for the case of pairwise relaxation dynamics in Section 4.1 and then relate this to the population level behaviour in Section 4.2. Motivated by our findings that ensuring only stability is insufficient, we then perform a convergence study in Section 4.3 where we compare first and second-order solvers.

### 4.1 Requiring numerical stability is not enough to prevent unphysical trajectories after cell division

The forward Euler method is the numerical scheme used most often in CBM implementations. Therefore, we started by studying the effect of the time step on the pairwise relaxation dynamics of cells after division, when simulated with this method. We use the same experimental setup as in Section 3.2.1, but now vary the time step Δ*t* used to numerically solve the ODE system as given in Equation (7).

We show the relaxation dynamics for the three force functions and different time steps in Figure 9. We observe three types of behaviours of the numerical solution depending on the time step size. Firstly, if the time step is too large, the movement of the cells centres in one time step can be so large so that cells immediately separate and remain far from each other for the duration of the simulation. This behaviour can be seen in Figure 9d for the cubic and the piecewise quadratic forces. The reason is that the centre-centre distance exceeds the adhesion threshold. Secondly, for smaller time steps, we can observe dynamics where the cells overshoot but adhere and we eventually recover the correct separation distance at equilibrium (Figure 9b for the cubic force and Figure 9c for the cubic and piecewise quadratic force). Numerically, this corresponds to a stable solution for a time step Δ*t* smaller than a threshold 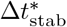. However, this behaviour leads to unphysical cell trajectories after division. Therefore, it is important that the time step is small enough to ensure monotone, smooth and accurate relaxation of daughter cells following the division of the mother cell. This third behaviour can be seen in Figure 9a for all three force functions, in Figure 9b for the piecewise quadratic and the GLS force and in Figure 9c only for the GLS force. From a numerical viewpoint, it can only be observed for time steps below a certain threshold 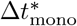. The exact thresholds for this critical time step are force function dependent.

**Figure 9:**
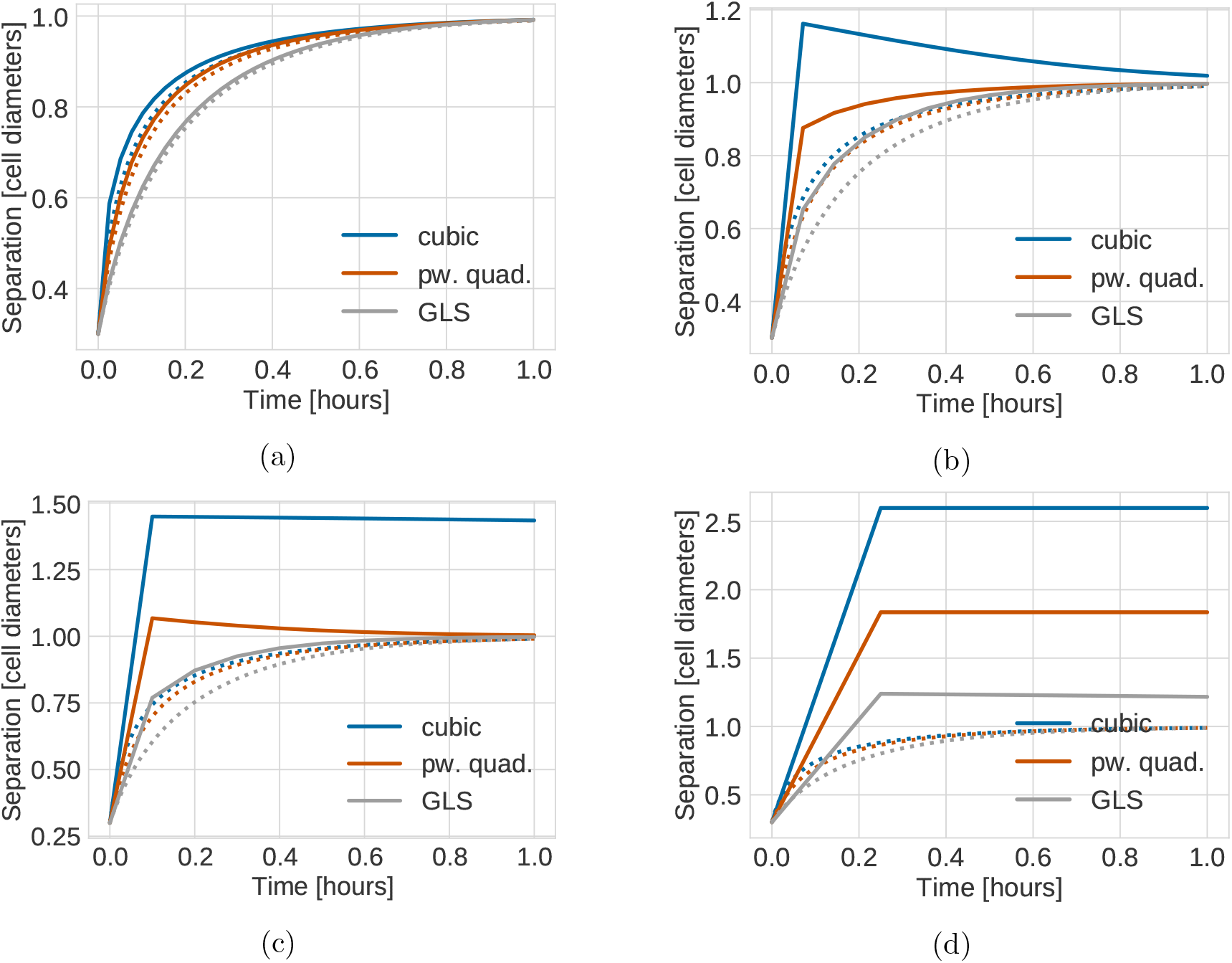
Stability of the numerical solution for the relaxation dynamics between daughter cells under different force functions. (a) Δ*t* = 0.025*h*, (b) Δ*t* = 0.075*h*, (c) Δ*t* = 0.1*h*, (d) Δ*t* = 0.2*h*. For reference, the dotted curves correspond to an accurate solution (less than 1% relative error) calculated using Δ*t* = 0.005*h*.

For the case of pairwise relaxation after cell division, as we consider here, the thresholds for stability and monotonicity can be calculated as

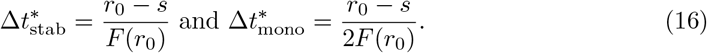

Note that the monotonicity threshold is exactly half the stability threshold. Since the force is negative and monotonically increasing for separation distances smaller than the rest length, i.e. *r* < *s*, the bound will be determined by the restriction on Δ*t* in the first step and hence will depend on the initial separation distance *r*_0_ between cells after division. 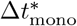 implies an upper limit on the error in the transient phase for pairwise relaxing cells in order to ensure physical division trajectories. When comparing different numerical schemes, this limit will be met for different 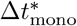. The error, and hence 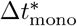, depends both on the relaxation time (stiffness) and the local truncation error of the scheme. The force function chosen directly affects the stiffness of the system. Table 4 lists the threshold values for the three force functions. When fitted to have the same relaxation time, the GLS force allows for larger time step sizes than the piecewise quadratic force. The cubic force function has the smallest, i.e. the most restrictive bounds.

**Table 4:** Table containing the monotonicity bounds 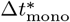 and stability bounds 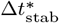 for the relaxation dynamics of pairwise overlapping cells for the different force functions (values given in hours), obtained by evaluating Equation (16) for *r*_0_ = 0.3 cell diameters and the force parameters according to Table 2.

| Force function | $\Delta t_{\text{mono}}^*$ | $\Delta t_{\text{stab}}^*$ |
| --- | --- | --- |
| Cubic | 0.0609 | 0.1218 |
| Pw. quad. | 0.0912 | 0.1823 |
| GLS | 0.1491 | 0.2982 |

In Figure 10 we studied the dependence of the stability and monotonicity bounds on the free parameters of the three force functions. We observed that the qualitative dependence of the bounds on the spring stiffness parameter agrees across all three forces. However, the parameter point corresponding to a consistent relaxation time of 1 hour is situated very differently on the curves, resulting in large quantitative differences in the threshold values for our parameterization.

**Figure 10:**
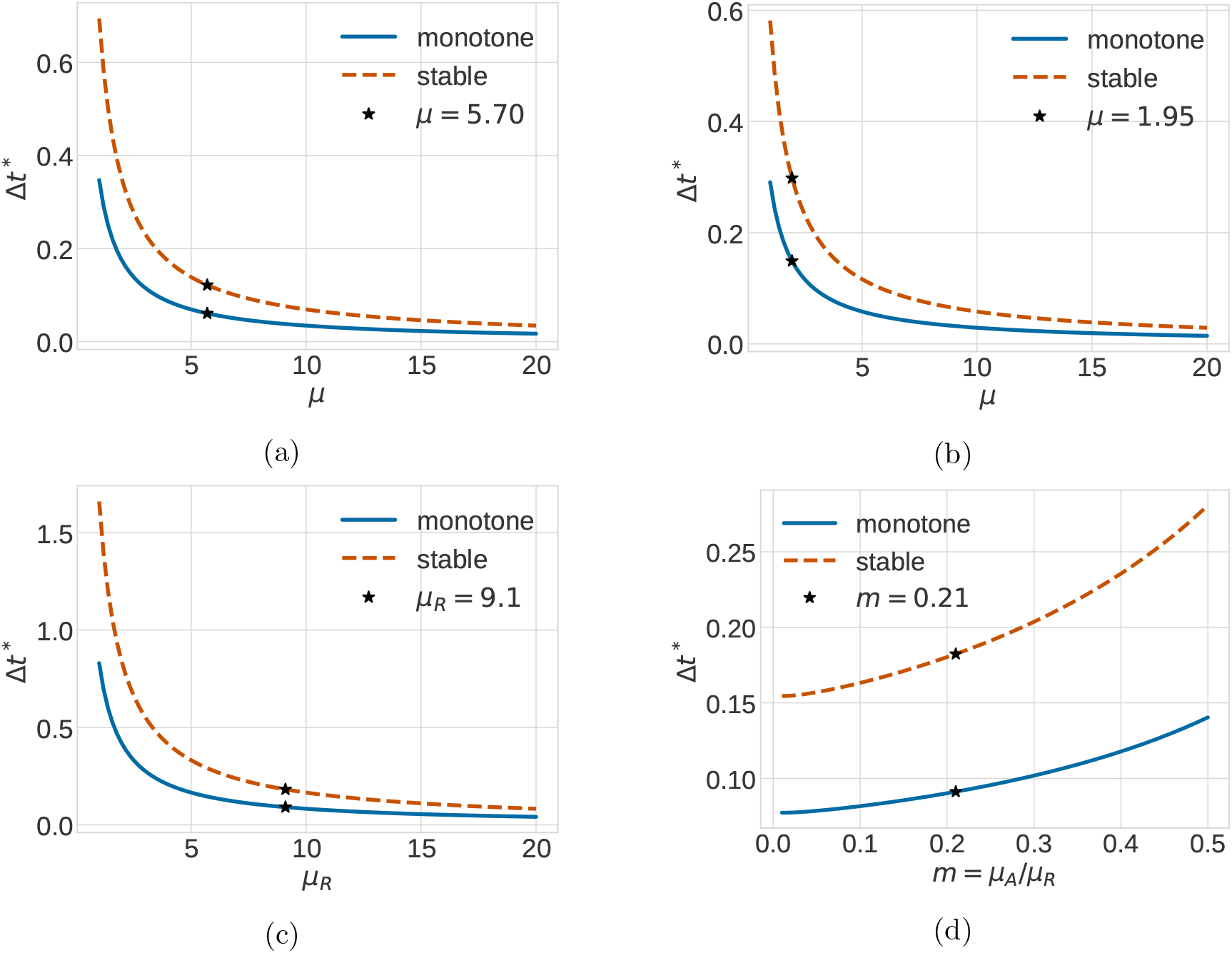
Stability and monotonicity bounds on Δ*t* given by Equation (16) as a function of force function parameters. The stars highlight the parameter values for which the force functions agree on a relaxation time of 1.0 hours.(a) cubic force, (b) GLS force, (c) piecewise quadratic force, fixed ratio *m* = 0.21, (d) piecewise quadratic force, fixed repulsive spring stiffness *μ_R_* = 9.1.

This points to another related, important practical consideration for a modeler, namely that the numerical properties for any scheme will depend on the physical parameters of the force function. When conducting e.g. parameter sweeps to study the sensitivity of some metric of interest, it is important to remember this since if the same time step is used and it is too large, one can easily end up in situations where one starts violating the monotonicity condition. Figure 11a quantifies this by plotting the monotonicity threshold 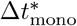 as a function of the spring stiffness parameter for the three force functions in a single plot. As can be seen, all force functions show the same qualitative behaviour as *μ* is varied. Since the spring stiffness *μ* scales inversely with the relaxation time *t*_0_ (compare with the discussion in Section 3.2.1), the dependence of 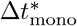 on the relaxation time is linear, as shown in Figure 11b.

**Figure 11:**
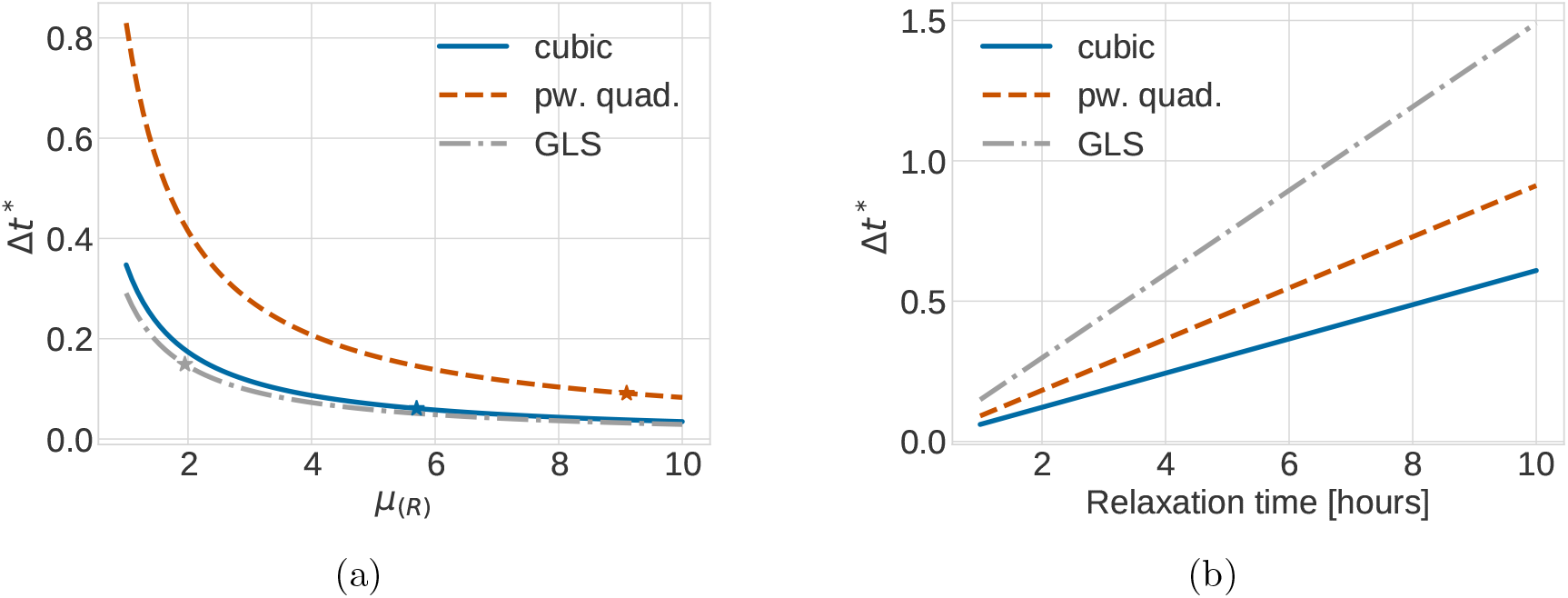
Comparison of the monotonicity bound 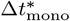 (c.f. Equation (16)) as a function of (a) the spring stiffness *μ*, respectively *μ_R_* and (b) the relaxation time for the different forces. The stars highlight the parameter values for which the forces agree on a relaxation time of 1.0 hours as given in Table 2. For the piecewise quadratic force the ratio was fixed as *m* = 0.21.

We can conclude that requiring stability of the numerical solution is insufficient for ensuring that cell trajectories are physically correct after division. Furthermore, we observed that the three forces put different restrictions on the time step size, with the GLS force allowing for the largest step sizes while still qualitatively correctly resolving the dynamics for daughter cells after division. We now move on to relate our findings for this case of pairwise interaction to the population level behaviour of a monolayer population under intense proliferation.

### 4.2 Too large step sizes can lead to geometrical differences at the population level

In the previous section, we studied the case of pairwise interactions between cells. These bounds will not hold exactly for more complex systems because the underlying ODE system to be solved is more involved. Similarly to Section 3 we here empirically relate the findings for the pairwise case to the case of simulating the positions of cells and the population radius of a monolayer population. Although it is unclear how to precisely relate the two cases mathematically, one can expect that not ensuring physicality of the trajectories of the daughter cells after cell division will have an impact on the population level behaviour.

The three upper rows of Figure 12 show an example realisation of a monolayer population relaxing to a mechanical equilibrium after simultaneous cell divisions for different time steps under the three force functions. The left most column corresponds to an accurate solution at *t*_end_ = 10 *h*, and the other two columns show the population for larger time step values at the same end time. We note that all chosen time steps lie within the stability bounds calculated for the pairwise relaxation case in the previous section, but not all lie within the monotonocity threshold (compare with Table 4). The random seed is fixed across time steps and force functions for this experiment, resulting in a consistent initial condition. This means that all observed differences in the geometrical configurations are purely due to the accuracy of the numerical solution and to potential differences in the force functions.

**Figure 12:**
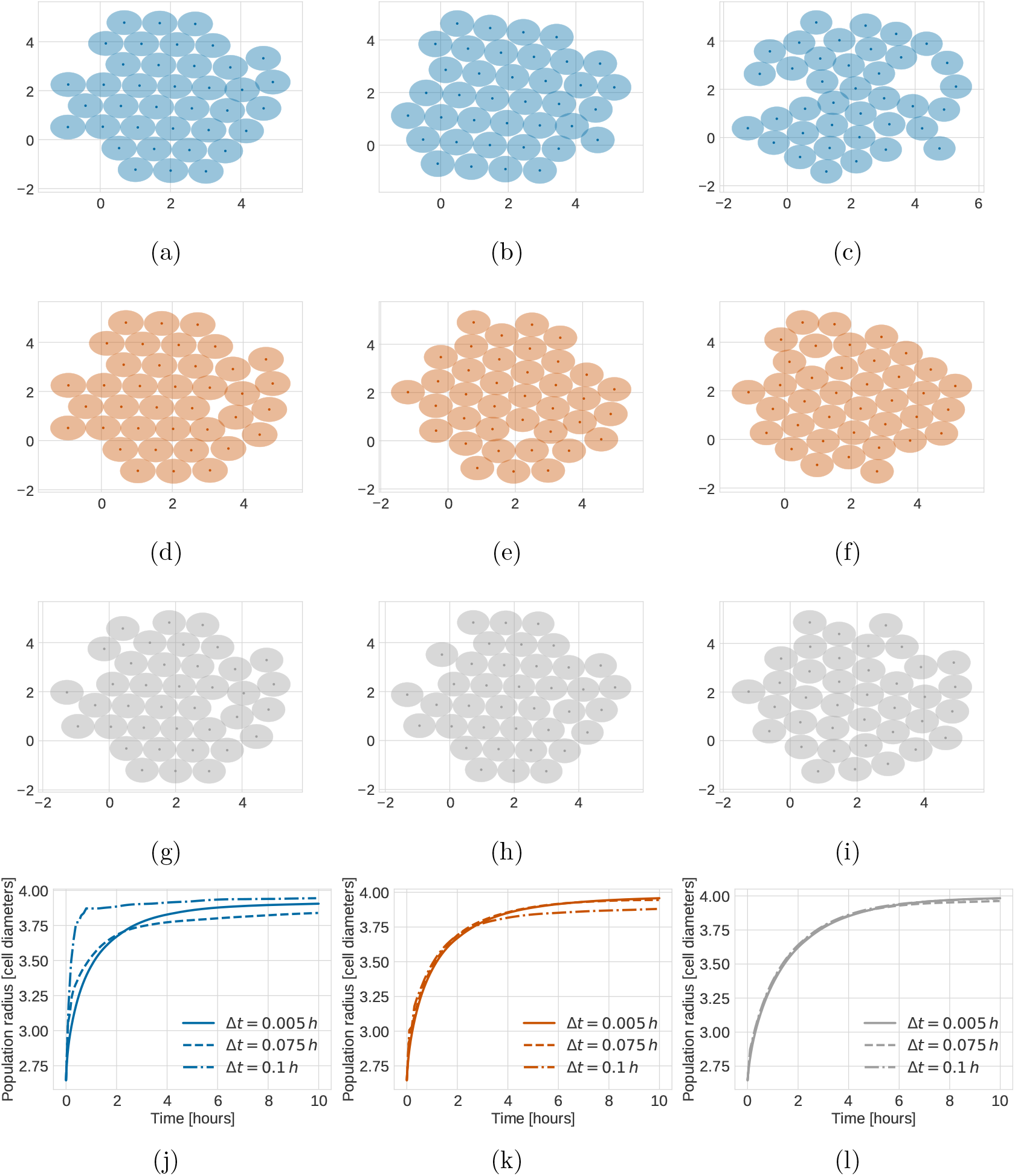
Impact of the time step size on the geometrical shape of a population and its radius. First row: 2D plots of a monolayer population simulated with the cubic force simulated for 10 hours with the forward Euler method and different time step sizes, (a) Δ*t* = 0.005 *h*, (b) Δ*t* = 0.075 *h* and (c) Δ*t* = 0.1 *h*. Second row: same setup simulated with the piecewise quadratic force. Third row: same setup simulated with the GLS force. All time step values lie within the stability region of the forward Euler method for all three force functions. The last row shows the radius of a monolayer population over time. Initially the population consisted of 19 cells. All cells divided simultaneously once at the beginning of the simulation. Then the population was allowed to relax to a mechanical equilibrium configuration. R esults a re a veraged o ver 1 0 simulation runs. The force function used was (j) the cubic force, (k) the piecewise quadratic force and (l) the GLS force.

As can be seen by comparing the simulations in the left most columns (Figures 12 a, d and g), we observe an almost identical positioning of the cells for the piecewise quadratic and GLS forces, with only slight differences from the cubic force for an accurately resolved simulation. To quantify this, the relative differences in centre positions between simulations using the GLS and the piecewise quadratic forces is 2.9% and between the GLS and cubic forces 3.1%. These simulations are computed with a small time step Δ*t* = 0.005 *h*, leading to a relative discretization error in centre positions of 0.5% for the cubic force and slightly less for the piecewise quadratic (*ϵ*_rel_ = 0.4%) and the GLS forces (*ϵ*_rel_ = 0.3%). This illustrates that the parameterization strategy we have chosen, i.e. to match the pairwise relaxation time, also results in very similar dynamics even on the level of centre positions for initial stages of a two-dimensional monolayer growth given an accurate enough numerical solution to the equations of motion.

On the other hand, we observe relatively large geometrical deviations from the well-resolved case for all three force functions as we increase the time step sizes, but like in the pairwise case, the sensitivity depends a lot on the force function (Figures 12 c, f and i). Consistent with the fact that it had the strictest threshold for dynamics of pairwise relaxing cells, these geometrical differences are biggest for the cubic force function (Figures 12 a–c), and again, the GLS force is most robust to variations in the time step (Figures 12 g–i) with the piecewise quadratic force falling in between (Figures 12 d–f). This is more easily seen in Figures 12 j–l where we show the population radius as a function of time for different step sizes for each of the force functions. Again, the radius is most sensitive to the time step under the cubic force.

In summary, the experiment in this section illustrates that the consequences on population level behaviour can be substantial if the time step is not chosen with care. Again, we emphasize that for an accurate numerical solution (leftmost column), simulations with all three force functions – if appropriately parameterized – result in geometrically very close solutions, whilst the sensitivity to the time step size differs substantially.

### 4.3 Convergence study for first- and second-order explicit methods

In order to quantify the effect of the time step, we conducted a convergence study for each of the three experimental setups we have considered until now: (i) relaxation of overlapping cells after cell division simulated until *T* = 1 *h*, (ii) pairwise dynamics of closely adhering cells, initially placed at a distance of 1.15 cell diameters and simulated until mechanical relaxation at *T* = 3 *h* and (iii) mechanical relaxation of two small proliferating monolayers of initially 19 and 37 cells simulated until *T* = 4 *h*. We considered the forward Euler method, the midpoint method and the Adams-Bashforth method. The forward Euler method — used in the majority of CBM software — is first-order, as confirmed by the convergence graphs. The midpoint and the Adam-Bashforth methods are commonly used second-order schemes, where the latter is also the default solver in the PhysiCell software [27].

Figure 13 shows the relative error in the numerical solution as a function of time step, force function and numerical scheme. For a discrete time step Δ*t* we computed the relative error as

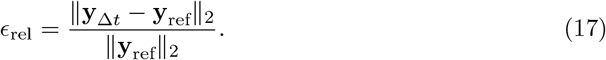

Here, **y**_ref_ is a gold standard reference solution computed with a very small time step for the respective force function. We used Δ*t*_ref_ = 1 × 10^−5^ *h* for case (i), Δ*t*_ref_ = 1 × 10^−4^ *h* (ii), and Δ*t*_ref_ = 5 × 10^−4^ *h* for case (iii). We interpolated the coarser solution **y**_Δ*t*_ down to the fine time grid used for **y**_ref_.

**Figure 13:**
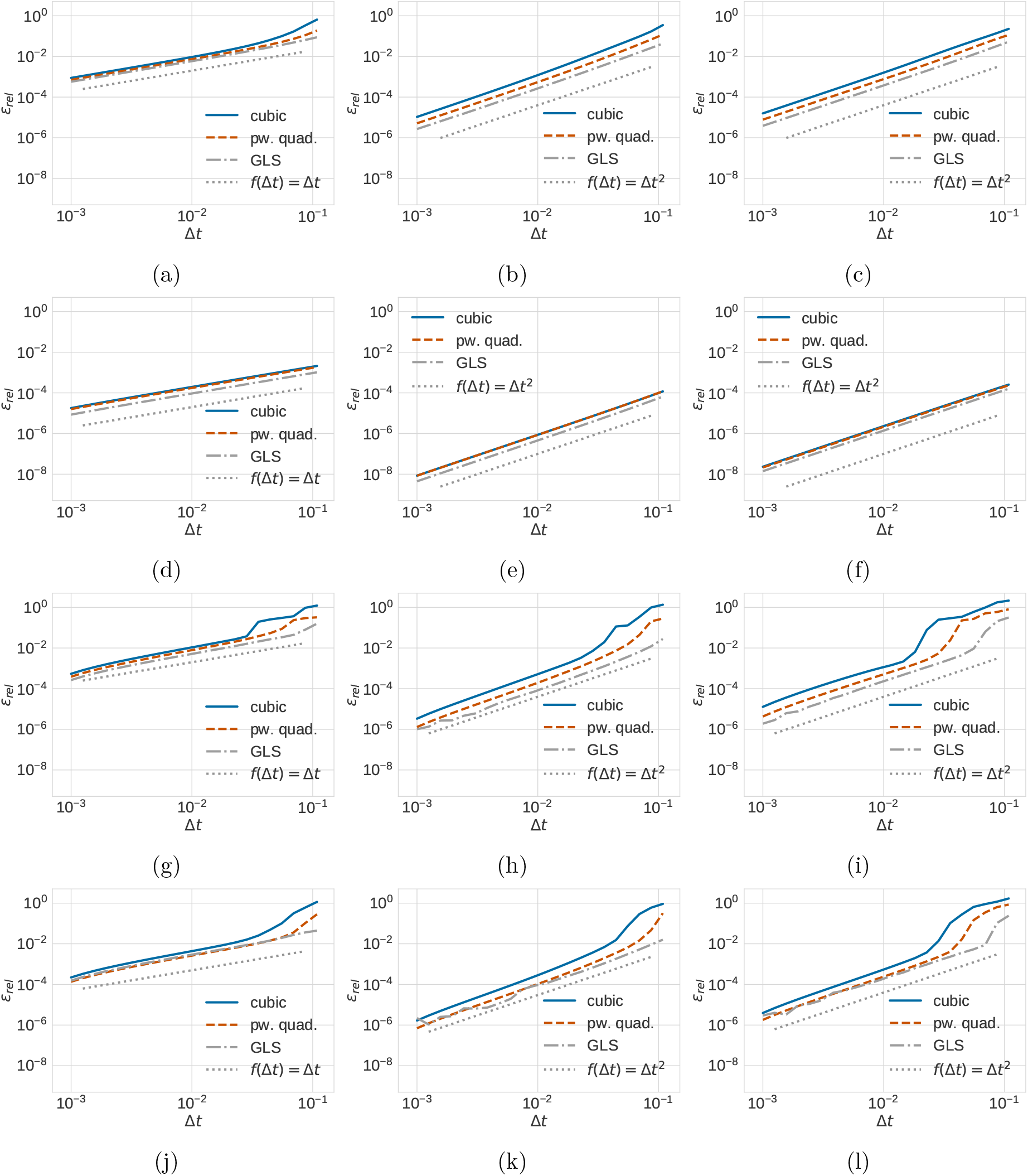
Impact of time step size on the relative error for the different combinations of forces and numerical solvers tested on the three model problems (i)-(iii). (a-c) Convergence study for the relaxation experiment (i), initial separation 0.3 cell diameters, (d-f) convergence study for adhering cells (ii), initial separation 1.15 cell diameters, (g-i) convergence study for a monolayer population of initially 19 cells and (j-l) convergence study for a monolayer population of initially 37 cells (iii). The numerical solver used was (a, d, g, j) the forward Euler method, (b, e, h, k) the midpoint method and (c, f, i, l) the Adams-Bashforth method.

As can be seen in Figure 13, all three tested schemes show the expected convergence rate — first-order for the forward Euler method and second-order for the midpoint and the Adams-Bashforth methods — for all three test cases, i.e. for the pairwise relaxation case (Figures 13 a–c), for relaxation in the adhesive regime (Figures 13 d–f) and for the centre positions in two monolayers of different sizes(Figures 13 g–i and 13 j–l), provided Δ*t* is chosen small enough. The errors are larger overall for the repulsive part than for the adhesive one. This means that if we choose a time step that accurately resolves the transient repulsive regime we do not need to worry about the adhesive regime. This holds for all solvers and all force functions.

From the convergence study, we can draw some conclusions about the computational efficiency associated with the numerical methods and force functions. As can be seen, both second-order methods allow for approximately 4–5× larger time steps than the forward Euler method at a relative error level of 1%, and up to approximately 10× at 0.1%. This is a substantial difference that is likely to lead to computational efficiency improvements in a state-of-the art implementation, even if two force function evaluations are needed at each time step rather than a single one for the forward Euler method. The midpoint method shows slightly better properties than the Adams-Bashforth method for large time steps. For this parameterization of the force functions, which results in close two-dimensional monolayer dynamics, the GLS force allows for up to 3–4× larger time steps with all solvers compared to the piecewise quadratic force, which in turn allows for 2× larger time steps than cubic. This is consistent with the observations in the previous section and is a consequence of the faster asymptotic drop-off in force for the cubic force (it behaves more rigidly than the piecewise quadratic force which in turn leads to more rigid cells than GLS). For the larger monolayer the GLS force seems to loose its advantage over the piecewise quadratic force.

In summary, assuming we want the relaxation dynamics after cell division resolved to a relative error of 1% in our simulation, the difference in time step between choosing the GLS force function and the midpoint method versus the cubic force and forward Euler method is an order of magnitude in our experiments. However, it will lead to almost identical simulations of the radius of a growing monolayer. This highlights the fact that substantial computational savings can be expected if force and numerical method are chosen carefully. We note here that there are of course other differences between the force functions that might affect a modelers choice, such as the degree of flexibility with which repulsion and adhesion can be chosen independently of each other. We also note that it is in general hard or impossible to *a priori* relate the error for the simple pairwise case to more complex summary statistics of simulations of large, complex systems. For this reason, a good recommendation for choosing the time step *a priori* is to choose it such that the error in pairwise relaxation is small, since it is likely that the case of two cells undergoing transient relaxation from a highly compressed state represents the worst-case behaviour in a complex tissue simulation since there are no balancing, opposite forces from surrounding cells.

From a pure numerical robustness point of view, the GLS force shows some advantages over the piecewise quadratic and the cubic force in our experiments, however here there are situations that need to be treated carefully. Recall that in our study in earlier sections, we had chosen its parameter *α* such that we assure a smooth drop-off to zero in force magnitude at the adhesive cut-off *r_A_*, see Equation (14) and Figure 3f. Figure 14 shows the convergence of the forward Euler, the midpoint and the Adams-Bashforth methods for the GLS force when we instead chose *α* = 5.0 independently of the spring stiffness parameter *μ* (the default choice in the Chaste software which uses the forward Euler method as a numerical solver). As can be seen, this choice of parameters leads to a loss of second-order convergence for both the midpoint and the Adams-Bashforth methods. This is because of the discontinuity in the force at the cutoff *r_A_*. This illustrates that in particular when considering the use of higher order methods, care needs to be taken to assure sufficient smoothness of the force function. The higher order of the method, the more smoothness is needed.

**Figure 14:**
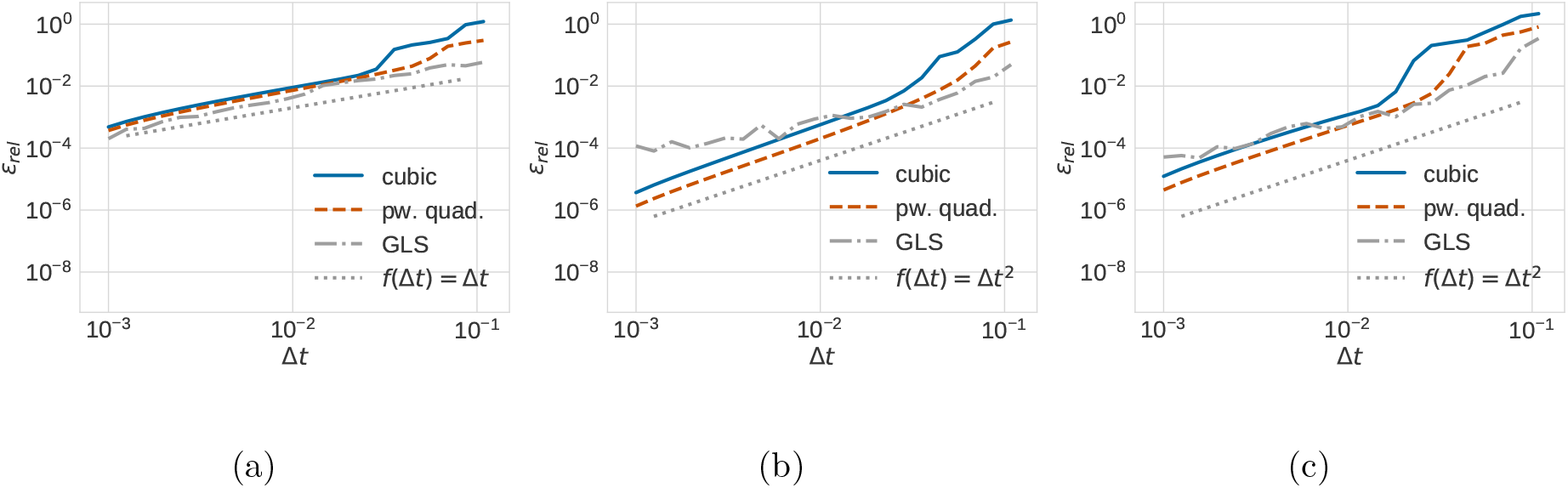
Convergence study for a monolayer population consisting initially of 19 cells. The parameter *α* of the GLS force was chosen independently of the spring stiffness *μ* as *α* = 5.0. The numerical solver used was (a) the forward Euler method, (b) the midpoint method and (c) the Adams-Bashforth method.

### 4.4 Comparison of execution times for first- and second-order explicit methods

To summarize our findings from the previous section, we should in general be able to expect computational efficiency gains of using a second-order solver such as the midpoint or the Adams-Bashforth method over the forward Euler method (unless we allow for quite large errors in the pairwise relaxation dynamics (around 10% or higher)). Their second-order convergence means that this advantage grows the more well resolved we want our simulation to be. To get a feel for the gain, we lastly compared execution times for first- and second-order solvers. Figure 15 shows the wall clock time for the simulation of a monolayer of 400 cells for the different combinations of forces and numerical solvers. As in our other experiments, all cells divided simultaneously at the beginning of the simulation. The time step sizes were chosen to satisfy a relative accuracy of *ϵ*_rel_ = 10^−3^ and are stated in Table 5 along with the number of steps of that size necessary to simulate the mechanical relaxation of the monolayer for 4 hours in-simulation time. We observe significant gains of using the second-order midpoint and Adams-Bashforth methods for all three force functions compared to using the forward Euler method. For example using the piecewise quadratic force and the midpoint method required less than 20 seconds, whereas using the same force and the forward Euler method took more than 40 seconds, i.e. double the time. Again, the cubic force performed slightly worse than the GLS and the piecewise quadratic forces, due to needing smaller time steps to resolve to the same accuracy (compare with Table 5). We stress that these gains can be expected to be even larger for a high-performance implementation.

**Figure 15:**
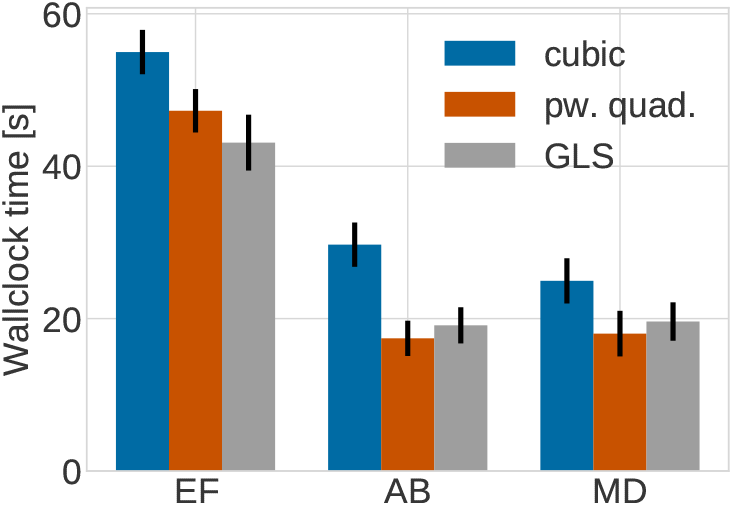
Execution time for a relaxing monolayer initially consisting of 400 cells for a relative accuracy of *ϵ*_rel_ = 10^−3^ for the different combinations of forces and numerical solvers. All cells divided at the beginning of the simulation. The monolayer then relaxed to mechanical equilibrium until time *t*_final_ = 4 *h*. EF - the forward Euler method, MD - the midpoint method, AB - the Adams-Bashforth method.

**Table 5:** Time step Δ*t* and number of steps *N* required for resolving the trajectories of a monolayer of 400 cells until time *t*_final_ = 4 *h* for a relative accuracy of *ϵ*_rel_ = 10^−3^ for the different combinations of forces and numerical solvers.

|  | Forward Euler | Midpoint | Adams-Bashforth |
| --- | --- | --- | --- |
| Cubic force | $\Delta t = 0.0050, N = 800$ | $\Delta t = 0.0238, N = 169$ | $\Delta t = 0.0191, N = 210$ |
| Piecewise quadratic force | $\Delta t = 0.0063, N = 635$ | $\Delta t = 0.0373, N = 108$ | $\Delta t = 0.0373, N = 108$ |
| GLS force | $\Delta t = 0.0122, N = 328$ | $\Delta t = 0.0582, N = 69$ | $\Delta t = 0.0582, N = 69$ |

## 5 Discussion and conclusion

In this paper we have systematically studied to what extent the choice of the pairwise interaction force in CBM OS simulations affects the numerical robustness and accuracy of simulations. We have also introduced an empirical way of calibrating different forces to give a similar population-level behaviour for two-dimensional monolayer growth.

The motivation for our study was twofold. Firstly, when engaging in modelling projects using popular open-source tools such as Chaste and Physicell, the question naturally arises to what extent the different default forces used will lead to any significant difference in the results. Secondly, given the assumption that the force functions are interchangeable from a modelling perspective, we were interested in whether their detailed form and implementation could have a significant impact on the simulation efficiency. To study these questions, we considered the simplest possible implementation of a centre-based model (as described in Section 2). There are many more physically realistic and intricate implementations of neighbourhood definitions, the cell cycle and cell proliferation. However, this simple formulation is still widely adopted since it is most computationally efficient. This is important for a CBM, since the target cell counts in simulations are in the millions or higher, at least in the case of high-performance and parallel implementations.

To be able to quantitatively compare numerical properties we focused on two model systems. The first was the pairwise relaxation dynamics following the placement of two cells with an overlap *r*_0_, as considered similarly in the supplementary information of the PhysiCell software [27]. The second was early stages of two-dimensional monolayer growth. By asking that all three force functions we considered should result in the same (or rather very close) population radius, we could parameterize all forces to agree qualitatively. Interestingly, matching the pairwise relaxation times resulted in almost the same population-level dynamics for two different monolayer sizes. Future work should check if this still holds true for spheroid growth in three dimensions. Based on this parameterization we could conclude that from a purely numerical efficiency point of view, there can be significant performance differences between the studied force functions if the objective is to simulate the same population-level metric.

Our study complements both the literature on comparisons of different types of CBMs [21, 38] and other careful evaluations of numerics for one particular force function [27]. One central issue we focused on is the possible overshoot for too large time steps for highly compressed cells right after cell division due to unphysically large force magnitudes. This problem is well-known in the CBM methods developer community and different ways of dealing with it have been suggested [23, 44]. Since the solution does not necessarily become numerically unstable and obviously wrong (see Figure 12), there is a real risk for misinterpretations of results, not only quantitatively (which would be less of an issue given the large uncertainties in the model and the many model assumptions), but also qualitatively. An example of this are the clear differences in the visual appearance of the geometry, for example a more loose structure with diffuse borders in Figure 12c) caused by time steps violating the monotonicity threshold. There is a risk for attributing this difference to the physical parameters of the model. We contributed here with a quantitive, numerical analysis of this problem for the model problem of two cells relaxing right after cell division. In particular, we show how to compute limits on the time step to ensure a monotone solution without overshoot and compare such limits for the three force functions under study. Again, we emphasize that even for the forward Euler method – the dominating scheme used in implementation to solve the functions of motion– these limits are accuracy-based and not based on the formal numerical stability of the solution (this bound is twice as large for the forward Euler method).

In comparing errors and convergence with two commonly used second-order schemes, we found it likely that second-order schemes, such as the Adams-Bashforth method used in PhysiCell [27] should provide a computational gain in high-performance implementations even if they require a more expensive update step. The gain becomes larger and larger the more accurately cell trajectories need to be resolved but outperforms the forward Euler method in our experiments even for modest accuracy requirements. However, as seen in Section 4.3, care needs to be taken with the GLS force to choose parameters to preserve second-order convergence for second-order schemes. In general, sufficient smoothness of the force is a requirement for higher-order convergence and would likely become an issue also for the piecewise quadratic force for e.g. Runge-Kutta schemes unless *n* is chosen to match the order of the scheme [27]. We also note here that the challenge is to resolve the transient solution right after cell division and while a higher-order scheme allows for larger time steps than the forward Euler method, another possible approach to improve efficiency is time step adaptivity [44, 23, 60]. Of the forces we considered, the GLS force used in Chaste [24] has the best numerical robustness properties and leads to the lowest discretization errors, both for first- and second-order solvers. The cubic force function was most sensitive to key to changes in numerical parameters and required overall smaller time steps than both the piecewise quadratic and the GLS forces.

It is not possible to make a firm statement about what combination of force function and solver to use as a default based only on our study, since there are both qualitative and quantitative aspects to that question. For example, from a qualitative point of view, the piecewise quadratic force used in Physicell [27] permits greater flexibility to vary the repulsive and adhesive spring stiffness parameters independently of each other, which is a very good feature for parametric studies. Other physics-based force functions not studied here such as the extended Hertz force [34] have the advantage of experimentally measurable parameters. The GLS force is the least flexible when it comes to the ability to be tuned to match dynamics of other force functions and it changes functional form in the adhesive regime.

Finally, there are many important considerations when choosing a force function, such as physical realism, modelling flexibility, ease of implementation, etc. However, given a specification of those requirements, we demonstrated here that the difference in time steps for a given accuracy could easily be as big as an order of magnitude for a highly resolved simulation purely dependent on the specifics of the force choice. This suggests that it would be possible to construct a force function in a way that it has good, or perhaps even optimal numerical properties. To do so, we would suggest to establish base-line dynamics using the physics-based JKR force function. A primary objective would then be to construct a force function that a) is considerably simpler but can be cheaply parameterized *a priori* to yield simulation results that match population-level dynamics of JKR quantitatively for a given set of parameters, and b) has as slow drop off of force magnitudes as possible for highly compressed cells (without distorting population-level metrics). More work could be done to evaluate the impact of the hysteresis effect of the JKR force (cells only adhere if they have been in contact before) on the population-level behaviour as this is something that none of the forces considered in our study can capture.

## Acknowledgments

We thank Per Lötstedt for productive discussions around the content of this article. We thank Jochen Kursawe for helpful comments on an earlier version of the manuscript. This work has received funding from the Swedish Research Council under grant 2015-03964 and from the eSSENCE strategic initiatives on eScience. The computations were performed on resources provided by SNIC through the Uppsala Multidisciplinary centre for Advanced Computational Science (UPPMAX) under Project SNIC 2019/8-227.

